# Hyaluronic Acid-Coated Melt Electrowritten Scaffolds Promote Myoblast Attachment, Alignment, and Differentiation

**DOI:** 10.1101/2025.03.06.641880

**Authors:** Alycia N. Galindo, Alyssa K. Chi, Ievgenii Liashenko, Kelly L. O’Neill, Chandler L. Asnes, Ruchi Sharma, Jenna D. Khachatourian, Armaan Hajarizadeh, Paul D. Dalton, Marian H. Hettiaratchi

## Abstract

**Purpose:** In muscle tissues, anisotropic cell alignment is essential for optimal muscle fiber development and function. Biomaterials for muscle tissue engineering must guide cellular alignment while supporting cell proliferation and myogenic differentiation.

**Methods:** Here, we describe the fabrication of a tissue-engineered construct consisting of a scaffold of aligned poly(ε-caprolactone) (PCL) microfibers coated in a dynamic covalent hydrazone crosslinked hyaluronic acid (HA) hydrogel to support myoblast attachment, myoblast alignment, and myotube formation. Norbornene modification of HA further enabled functionalization with fibronectin-derived arginine-glycine-aspartic acid (RGD) peptide. Scaffolds were fabricated using melt electrowriting (MEW), a three-dimensional (3D)-printing technique that uses stabilization of fluid columns to produce precisely aligned polymeric microfibers. We evaluated C2C12 mouse skeletal myoblasts cultured on non-coated, HA-coated, and HA-RGD-coated MEW scaffolds with fiber diameters of 10 µm, 20 µm, and 30 µm using immunocytochemistry and creatine kinase activity assays. We further evaluated the mechanical properties of 20 µm fiber scaffolds and their effect on myogenic gene expression and alpha-actinin protein expression of C2C12 myoblasts undergoing differentiation.

**Results:** HA-coated and HA-RGD-coated scaffolds increased attachment of C2C12 myoblasts on all fiber diameters compared to non-coated scaffolds, with HA-RGD-coated scaffolds demonstrating the highest cell attachment. All scaffolds supported cellular alignment along the fibers. Cells differentiated on scaffolds showed anisotropic alignment with increased myotube formation on HA-coated and HA-RGD-coated scaffolds as demonstrated by myosin heavy chain (MHC) staining and by the presence of striations on HA-coated scaffolds visualized with alpha-actinin staining. Increased creatine kinase activity and myogenic gene expression on day 5 further indicated myotube formation on all scaffolds, with HA-coated scaffolds significantly increasing the expression of several key myogenic markers.

**Conclusion:** This unique combination of tunable biophysical and biochemical cues enables the creation of a biomimetic tissue engineered scaffold, providing a platform for new therapeutic approaches for muscle regeneration.

## Introduction

Volumetric muscle loss (VML), characterized by extensive muscle loss of greater than 20% of its original volume, can be caused by trauma, chronic disease, or surgical resection.^1,2^ Such muscle injuries typically have limited regenerative capacity and cannot heal without additional intervention, often leading to fibrosis, fatty tissue infiltration, and impaired function, decreasing a patient’s quality of life.^3–5^ Muscle tissue has a complex and hierarchical architecture of aligned fascicles enclosed by the epimysium, the outermost layer of muscle.^6,7^ Within fascicles are bundles of muscle fibers that are surrounded by the perimysium. Individual muscle fibers within a fascicle are surrounded by endomysium. Muscle fibers, also known as myofibers, are multinucleated muscle cells. Within muscle fibers are myofibrils, which are responsible for muscle contractility and are made from repeating units of sarcomeres that contain thick myosin and thin actin filaments. VML destroys the natural muscle ultrastructure, preventing the muscle from performing essential tasks like contraction and force generation.^4^ Current therapeutic approaches for treating VML often require multiple interventions, including surgical procedures such as autologous muscle transfer,^8^ rehabilitation,^9^ and long-term management of associated complications.^10,11^ However, these approaches often fail to fully restore muscle function due to the difficulty of achieving aligned tissue,^12,13^ and restoring this hierarchical structure is considered an important step for the repair of functional muscle tissue following VML.^13,14^ Furthermore, the use of muscle autografts and allografts is constrained by the scarcity of available tissue and the potential for donor-site complications, underscoring the urgent need for alternative solutions.^15^

To address the shortcomings of current therapeutic approaches for treating VML, scaffold-based tissue engineering has emerged as a potential therapeutic strategy for stimulating the regeneration of functional, mature muscle tissue after VML. Strategies that combine three-dimensional (3D) scaffolds,^16,17^ cell-based therapies,^18^ and biochemical cues,^19,20^ have shown promise in treating VML. For example, aligned scaffolds have been fabricated using 3D printing of fiber-based scaffolds or hydrogels,^21^ injection molding or casting,^22^ microfluidics,^23^ unidirectional freeze drying or ice templating,^24^ and decellularized tissues.^25^ Fiber-based scaffolds have been fabricated from synthetic polymers such as polylactic acid (PLA), poly (lactic-co-glycolic acid) (PLGA),^26^ and poly(ε-caprolactone) (PCL)^27^ with 3D printing techniques such as solution electrospinning and have been used to align myoblasts with a tissue-engineered scaffold and promote their differentiation. These scaffolds may further be tailored to replicate the extracellular matrix (ECM) of native muscle tissue, generating an enhanced microenvironment for myoblast adhesion, migration, and differentiation.^15,28^ Scaffolds fabricated from decellularized ECM from native muscle tissue are promising for VML treatment because they retain the biochemical signals and aligned structure of muscle tissue, which can facilitate myoblast alignment and fusion; however, it is often difficult to tune the properties of decellularized ECM scaffolds without losing their bioactivity.^29,30^ Similarly, naturally-derived biomaterials such as collagen, gelatin, silk, fibrin, and alginate can be fabricated into aligned materials to promote myoblast alignment and fusion.^31,32^ Hydrogels made from type I collagen,^33,34^ poly(ethylene glycol) diacrylate (PEG-DA),^20^ and gelatin methacryloyl^35^ have also shown promise in supporting the development of tissue-engineered muscle constructs, as they mimic the hydrated environment of muscle tissue and have the added benefit of supporting vascularization and innervation within engineered tissues.^20,33–35^ However, challenges remain with synthetic fiber-based approaches that may suffer from ineffective integration with the host tissue and limited biochemical cues to support cell adhesion, proliferation, and differentiation, while hydrogel-based approaches have demonstrated a limited ability to support aligned muscle architecture and long-term tissue stability. Therefore, integrating synthetic fibers with hydrogels could be a promising approach for integrating complementary biochemical and biophysical cues, thereby addressing the weaknesses of each approach when used alone.

Incorporating biochemical cues into scaffolds enhances muscle tissue regeneration by providing essential signals that guide cellular behavior. These cues may include cells, electroconductive materials that improve cellular material interactions, growth factors that stimulate cellular activity, and peptide motifs that facilitate cell attachment.^36–39^ By mimicking the body’s natural ECM, these scaffolds offer a promising solution to accelerate muscle tissue repair and minimize tissue fibrosis.^40^ A variety of ECM-derived peptides, such as fibronectin, laminin, and elastin, have been used in tissue-engineered scaffolds to stimulate myoblast proliferation, highlighting the importance of cell-ECM interactions for driving muscle tissue regeneration.^20,41,42^ The fibronectin-derived peptide RGD has been previously shown to support muscle satellite cell survival, attachment, and myogenic differentiation within polyethylene glycol maleimide (PEG-MAL) hydrogels.^43^ However, achieving robust muscle tissue formation using bioactive scaffolds is still a challenge, and the ideal combination of biophysical and biochemical cues to create a tissue-engineered scaffold that mimics functional muscle tissue has yet to be determined.

To address these challenges, we have created a composite tissue-engineered scaffold that combines the alignment properties of a synthetic fiber-based scaffold with the cell-supportive microenvironment of ECM-derived hydrogels to promote the attachment, alignment, and myogenic differentiation of skeletal myoblasts with a unique combination of biophysical and biochemical cues. We used melt electrowriting (MEW), a 3D-printing technique that uses electrohydrodynamics to precisely deposit multiple layers of microfibers onto a collector, to produce aligned scaffolds.^44,45^ To promote C2C12 murine skeletal myoblast attachment, we embedded MEW scaffolds within hydrazone-crosslinked hyaluronic acid (HA) hydrogels that have been previously characterized by our lab and others.^46–48^ HA can directly interact with myoblasts through the CD44 and RHAMM cell receptors, promoting cell migration, adhesion, and proliferation.^49^ We modified HA with norbornene functional groups for the chemical conjugation of RGD to further promote myoblast attachment and alignment. We seeded C2C12 myoblasts on non-coated, HA-coated, HA-RGD-coated PCL microfiber scaffolds and evaluated their attachment to the scaffold, alignment along PCL microfibers, and differentiation into myotubes. We hypothesized that both HA and HA-RGD-coated scaffolds would increase myoblast attachment compared to non-coated MEW scaffolds, and HA-RGD-coated scaffolds would result in the highest cellular attachment and enhanced myogenic differentiation. We evaluated myotube formation through myosin heavy chain staining and creatine kinase activity assays over time. Overall, this HA-coated composite scaffold presents a unique combination of biophysical and biochemical cues that promote myoblast attachment, alignment, and differentiation and has the potential to be used as a tissue engineering scaffold to accelerate functional muscle regeneration.

## Methods and Materials

### Oxidized Hyaluronic Acid (HA-Ox)

As previously described, sodium hyaluronate (HA, 100 kDa) (Lifecore Biomedical LLC, Chaska, MN) was oxidized to reveal aldehyde groups.^46^ Briefly, 2% w/v HA (1g, 0.00264 mol) was dissolved in double distilled water (ddH_2_O) in a round bottom flask. The flask was covered with foil, and 0.6 molar equivalents of sodium periodate (NaIO_4_, 338.61 mg, 0.00158 mol) (Sigma Aldrich, St. Louis, MO) were added to the reaction. The reaction was stirred for 4 hours in the dark before being quenched with 5 molar equivalents of propylene glycol (VWR Chemicals, Radnor, PA) (1 g, 0.0132 mol). The solution was then dialyzed against ddH_2_O for 2 days before being filtered, frozen at -80°C, and lyophilized.

### Hyaluronic Acid-Norbornene (HA-Nor)

HA was functionalized with norbornene as previously described.^46,50,51^ An intermediate product, HA-tetrabutylammonium (HA-TBA) was first synthesized. 2% w/v HA (1 g, 0.00264 mol) was dissolved in ddH_2_O in a round bottom flask. AmberLite MB ion exchange resin (3 g, 0.00962 mol) (Sigma Aldrich) was added to the flask and stirred for 5 hours. Vacuum filtration was then used to remove the resin. The solution was collected and titrated with tetrabutylammonium-hydroxide (TBA-OH) (Sigma Aldrich) to pH 7. The solution was then filtered, frozen at -80°C, and lyophilized. HA-TBA (1 g, 0.00157 mol) was dissolved in anhydrous dimethylsulfoxide (DMSO) (Millipore Sigma, Burlington, MA) at 2% w/v in a round bottom flask, which was submerged in an oil bath at 45°C and purged with nitrogen gas. 3 molar equivalents of 5-norbornene-2-carboxylic acid (Nor-CHOO, 0.803 g, 0.00581 mol) (Sigma Aldrich) and 1.5 molar equivalents of 4-(dimethylamino)pyridine (DMAP, 0. 287 g, 0.00235 mol) (Sigma Aldrich) were stirred into the flask until completely dissolved. Lastly, 0.4 molar equivalents of di-tert-butyl dicarbonate (Boc_2_O, 0.137 g, 0.000627 mol) (Sigma Aldrich) were added to the flask, and the reaction was stirred under nitrogen for 20 hours. The reaction was quenched with an equal volume of cold ddH_2_O and dialyzed against ddH_2_O for 3 days. After dialysis, the solution was collected in a beaker and sodium chloride (VWR Chemicals) (1 g/100 mL of solution) was added. Acetone was added to the beaker (1 L/100 mL of solution) to precipitate the product. The precipitate was then collected by centrifugation and redissolved in ddH_2_O. The HA-Nor solution was then filtered, frozen, and lyophilized.

### Hyaluronic Acid-Norbornene-Adipic Acid Dihydrazide (HA-Nor-ADH)

1% w/v HA-Nor (0.125 g, 0.000240 mol) was dissolved in ddH_2_O, and 10 molar equivalents of adipic acid dihydrazide (ADH) Spectrum (Chemical MFG Corp, Gardena, CA) (0.418 g, 0.00240 mol) and a one molar equivalent of hydroxybenzotriazole (HOBt) (Chem Impex, Wood Dale, IL) (0.324 g, 0.000240 mol) were added to the reaction. The pH was adjusted to 4.75 using NaOH/HCl, and one molar equivalent of 1-ethyl-3-(3-dimethylaminopropyl)carbodiimide (EDC) (G Biosciences, St. Louis, MO) (0.0460 g, 0.000240 mol) was added to the solution. The pH was titrated to 4.75 and monitored for 2-4 hours until stabilized. The reaction was then stirred for 2 days before being dialyzed against 0.1 M sodium chloride for 2 days followed by ddH_2_O for an additional 2 days. The solution was then filtered, frozen at -80 °C, and lyophilized.

### Determination of Degree of Modification (DOM)

As previously described, the degree of modification of HA-Nor and HA-ADH-Nor was quantified using proton nuclear magnetic resonance spectroscopy (^1^H-NMR, 500Hz, Bruker, USA).^46^ The DOM for HA-Nor was calculated by integrating the vinyl peaks on the norbornene functional group from 5.7-6.3 ppm and normalizing to the n-acetyl methyl group on HA from 1.8-2 ppm (3H).^51^ For HA-Nor-ADH, the aliphatic chain peaks from 2.5-2.1 ppm and 1.7-1.44 ppm (8H) were integrated and normalized to the n-acetyl methyl group peaks on HA from 1.8-2 ppm (3H).^51^ A representative ^1^H-NMR spectrum can be found in the supplemental information **(Fig. S1)**.

To determine the DOM of HA-Ox, a hydroxylamine hydrochloride titration was performed by dissolving 100 mg of oxidized polymer in 20 mL of 0.25 N hydroxylamine hydrochloride containing 0.05% w/v methyl orange reagent for 2 hours.^46,52,53^ The pH of the solution was monitored and titrated with 0.1 M NaOH until the pH indicator changed from red to yellow at the end point pH of 4. **Equation 1** was used to calculate the DOM or the percentage of HA monomers that contained aldehydes.

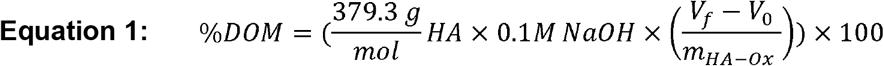

where 379.3 g/mol is the monomeric molecular weight of HA, 0.1 M is the molarity of NaOH used to adjust the solution pH, V_f_ is the final volume of NaOH in the burette after titration recorded in liters, V_0_ is the initial volume of NaOH in the burette before titration recorded in liters, and m_HA-Ox_ is the mass in g of HA-Ox used in the titration.

### RGD Conjugation to HA-Nor-ADH (HA-RGD)

2% w/v HA-Nor-ADH (10 mg, 0.0146 mmol) was dissolved in 500 µL of PBS containing 0.05% w/v of Irgacure 2959 (Sigma Aldrich) and 2500 nmol of RGD peptide with a cysteine at the N terminus (CGRGDSG) (GenScript, Piscataway, NJ) with the goal of conjugating 2.5 mM of RGD per 25 µL HA-Nor-ADH-RGD. The solution was exposed to UV light at a wavelength of 365 nm for 1 minute to perform a thiol-ene reaction between HA-Nor-ADH and RGD-cysteine.

The solution was dialyzed against ddH_2_O for 1 day before being filtered, frozen at -80°C, and lyophilized. Ellman’s reagent (Fisher Scientific, Waltham, MA) was used to detect free thiols and quantify the concentration of RGD conjugated to the HA-Nor-ADH through the disappearance of thiols after the photoinitiated reaction. The Ellman’s reagent consisted of 2 mM of 5,5-dithio-bis-(2-nitrobenzoic acid) (DTNB) and 50 mM sodium acetate in 1 M tris(hydroxymethyl)aminomethane (TRIS) buffer. Samples of the reaction before and after UV exposure were diluted to fall within the range of a standard curve of serially diluted RGD peptide. 50 µL of each standard and sample were added to a 96-well plate and mixed with 250 µL of Ellman’s reagent. The plate was developed at room temperature for 5 minutes prior to reading absorbance at 412 nm on a microplate reader (BioTek Synergy Neo2).

### Fabrication of Melt Electrowritten (MEW) Scaffolds

Three flat scaffold designs were fabricated from medical-grade PCL (PURAC PC12, Corbion Inc.) using a custom-built MEW printer.^54^ PCL was heated to 75°C and extruded through an electrically grounded 22 gauge nozzle protruding 1 mm from the printhead. The collector distance was fixed to 3.5 mm, and scaffolds were printed onto 1 mm thick glass microscopy slides resting on a metal collector. Three sets of parameters (including collector voltage, pressure delivered to molten polymer, and translation speed) were optimized to print fibers with nominal diameters of 10 µm, 20 µm, and 30 µm. Printing parameters together with corresponding scaffold designs are detailed in **Table 1**.

**Table 1.**
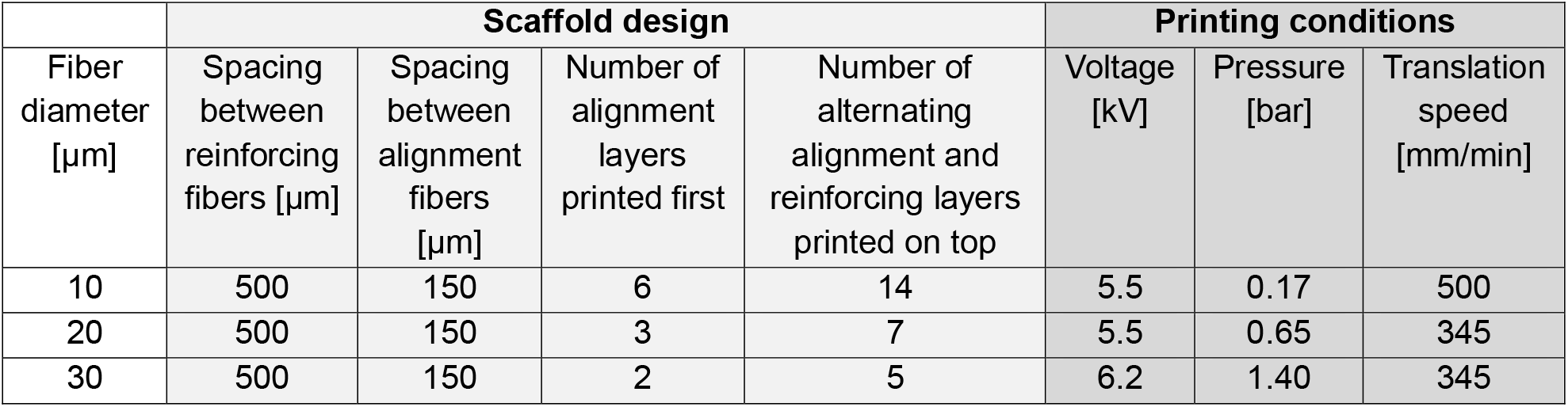

Square scaffolds (7x7 mm) consisting of two sets of perpendicular alternating fibers were printed. The first set of fibers intended for myoblast alignment were spaced 150 µm apart. The second set of fibers intended as reinforcement fibers ran perpendicular to the myoblast alignment fibers and were spaced 500 µm apart. The numbers of alignment fibers and perpendicular reinforcement fibers printed for each design are detailed in **Table 1**. The total number of layers for each fiber diameter was selected to provide a consistent scaffold height of approximately 200 µm across the different designs. Scaffolds printed with 10 µm fibers were additionally reinforced on the edges to enable handling through a buckling fiber rim design previously used to improve scaffold handling and printed at 1 mm/min translation speed.^55^

### Etching and Sterilization of MEW Scaffolds

Scaffolds were etched in a biosafety cabinet with 0.1 M NaOH for 30 minutes before being washed 3 times with phosphate-buffered saline (PBS). Etched scaffolds were disinfected in 70% v/v ethanol for 15 minutes before being washed three times with PBS. Scaffolds were stored in 10% v/v fetal bovine serum (FBS) (R&D Biosystems, Minneapolis, MN) in Dulbecco’s modified eagle medium (DMEM) (Cytiva, Marlborough, MA) (“high serum medium”) at 4°C overnight before use.

### HA Coating of MEW Scaffolds

HA-coated scaffolds were fabricated by adding 25 µL of either HA-Nor-ADH or HA-Nor-ADH-RGD to the bottom of an 8-chamber well slide (Fisher Scientific), followed by the scaffold, and then addition of 25 µL of HA-Ox. Coated scaffolds were agitated, and the polymers were allowed to crosslink for 30 minutes to create a thin HA hydrogel coating on the MEW scaffold.

### Scanning Electron Microscopy (SEM) Imaging and Fiber Measurements of MEW Scaffolds

HA-coated MEW scaffolds were prepared using a freeze-drying process for imaging with scanning electron microscopy (SEM). Freshly prepared HA-coated scaffolds were gently placed into a glass vial and suspended by a support on the edges. Samples were frozen at -80°C for 4 hours and then lyophilized overnight. Non-coated and HA-coated samples were then sputter-coated with 7 nm of titanium and imaged with an SEM Everhart–Thornley detector (ETD) (ThermoFisher Apreo 2, USA). Fiber diameter was quantified using ImageJ (FIJI, USA) **(Figure S2)**.

### C2C12 Myoblast Culture

C2C12 murine skeletal myoblasts (ATCC, Manassas, VA) were seeded at 2500 cells/cm^2^ in T-75 flasks with high serum medium at 5% CO_2_ and 37°C. C2C12 were passaged at 70-80% confluency following standard protocols before use in further experiments.^56^

### Myoblast Culture on Scaffolds

To evaluate myoblast attachment and proliferation, C2C12 cells between passages 7-9 were seeded on top of scaffolds at a density of approximately 57,000 cells/cm^2^ in an 8-chamber slide and allowed to attach to the scaffolds for 2 hours in a low volume (100 µL) of high serum medium. The remaining volume (300 µL) of high serum medium was added, and the cells were allowed to proliferate for 24 hours. The media was then exchanged to 1% v/v FBS in DMEM (“low serum medium”) for 3 days, after which the cells were fixed with 4% v/v paraformaldehyde (Fisher Scientific) in PBS for immunocytochemistry.

To evaluate myotube formation, C2C12 cells between passages 7-9 were seeded on top of scaffolds at a density of approximately 14,000 cells/cm^2^ in an 8-chamber well slide and allowed to attach for 2 hours in a low volume of high serum media before the remaining media was added. The cells were allowed to proliferate for 24 hours before the media was exchanged to differentiation media consisting of 2% v/v horse serum (Sigma Aldrich) in DMEM. Cells were differentiated for 7 days. The differentiation media was exchanged every 3 days. Media were collected at 1, 5, or 7 days to evaluate CK activity. After 7 days, cells were fixed with 4% v/v paraformaldehyde for immunocytochemistry.

### Immunocytochemistry

All immunocytochemistry steps were performed at room temperature in the dark. For F-actin staining, cells were permeabilized with 0.5% v/v Triton-X 100 (Sigma Aldrich) for 15 minutes before washing three times with 200 µL of PBS. 200 µL of Alexa Fluor™ 647 phalloidin (5 U/mL) (Biotium, Fremont, CA) were added to the well and incubated for 20 minutes. The wells were then washed three times with PBS, incubated with 1 mg/mL of 4′,6-diamidino-2-phenylindole (DAPI) (Biotium) for 5 minutes, and washed three times with PBS again.

For myosin heavy chain staining, cells were permeabilized with 0.5% w/v Triton-X 100 for 15 minutes at room temperature and washed three times with PBS. Cells were blocked with 1% w/v bovine serum albumin (BSA) (Fisher Scientific), 22.5 mg/mL glycine (TCI Chemicals, Tokyo, Japan), and 0.1% v/v Tween-20 (Acros Organics, Geel, Belgium) in PBS overnight with agitation at 250 rpm at 4°C. The next day, the cells were washed three times with PBS, anti-fast myosin skeletal heavy chain antibody (ab91506, Abcam, Cambridge, United Kingdom) diluted 1:500 times in 0.1% w/v BSA in PBS was added to the wells, and the cells were incubated overnight with agitation at 250 rpm at 4°C. The following day, cells were washed with 0.5% v/v Tween-20 in PBS three times for 20 minutes each with agitation at 250 rpm at 4 °C. After washing, goat anti-rabbit IgG secondary antibody Alexa Fluor™ 488 (Invitrogen, Waltham, MA) diluted 1:200 in 0.1% w/v BSA in PBS was added to the wells and incubated for 1 hour with agitation at 250 rpm at 4°C, then washed three times with PBS. Lastly, 1 mg/mL of DAPI was added to the well for 5 minutes at room temperature and then washed three times with PBS. Cells were stored in PBS for imaging.

Alpha-actinin staining was performed similarly to myosin heavy chain staining but using anti-alpha-actinin primary antibody (A7732, Sigma Aldrich, 1:67 dilution) and goat anti-mouse IgG secondary antibody Alexa Fluor™ 647 (Invitrogen, 1:1000 dilution) with similar incubation times and washing steps.

### Imaging of Scaffolds

C2C12 myoblasts were imaged as z-stacks (0.4 or 2.5 µm slices) using a CSU-W1 SoRa Spinning Disk confocal microscope (Nikon, USA). The average z-stacks were 68 µm for non-coated scaffolds, 110 µm for HA-coated scaffolds, and 129 µm for HA-RGD-coated scaffolds. Images were analyzed using IMARIS imaging software version 9.5. The Rolling Ball Algorithm was applied to each image to correct uneven backgrounds. The following background subtraction of fluorescent signal was applied to each stained channel (DAPI: 3500, Cy5: 3950, FITC: 3300) based on pixel size of the image.

### Nuclei Count Analysis

The “surfaces” analysis in IMARIS was used to quantify the number of nuclei per scaffold. The background subtraction width was set to 9 μm to remove any background based on the width of the nuclei. Seed point and surface filters were added to enhance nuclei count accuracy.

### Directionality Analysis

The directionality of myosin heavy chain staining was analyzed using the “cells” analysis in IMARIS. In the DAPI channel, the nucleus diameter (15 µm) and the average threshold intensity (327) were set. In the FITC channel, the average cell threshold intensity (173) and split by seed points (10 µm) were analyzed. The angle of the scaffolds was converted to degrees, and then the arctangent of “Cell Ellipsoid Axis X” and “Cell Ellipsoid Axis Y” was calculated and divided by two to get values between -90 to 90 degrees.

The corner points of two scaffold boxes were identified to determine the angle of the scaffold for directionality correction. The absolute value of the arctangent of the difference between the top and the bottom of the scaffold was determined and averaged for each scaffold. If the cell ellipsoid value was less than zero, the scaffold value was added to it. If the cell ellipsoid value was greater than zero, the scaffold value was subtracted from it.

### Creatine Kinase (CK) Activity Assay

Creatine kinase (CK) activity was measured using a Colorimetric CK Activity Assay kit (Abcam, Cambridge, United Kingdom) according to the manufacturer’s instructions. After Day 1, 5, and 7 of differentiation, cell culture medium was carefully aspirated from the cell culture chambers containing the scaffolds, transferred to sterile 1.5 mL microcentrifuge tubes, and stored at -80°C. Media samples (10 µL) were thawed and added to 96-well plates followed by the addition of a reaction mixture that included CK assay buffer, CK enzyme, CK developer solution, adenosine triphosphate (ATP), and creatine (CK substrate). The plate was incubated at 37°C for 40 minutes, and absorbance at 450 nm was measured using a microplate reader (BioTek Synergy Neo2). CK activity was normalized to protein concentration determined by a bicinchoninic acid (BCA) assay (Fisher Scientific). All samples were processed in technical triplicates. A negative control (cell culture medium from wells without cells) and a positive control (standard provided in the kit) were included in the assay.

### Mechanical Properties

The stiffness of non-coated MEW scaffolds was measured using the tension module on the RSA-G2 Solids Analyzer (TA Instruments, New Castle, DE). Scaffolds were attached at the top and bottom along the alignment fibers using tension clamps and allowed to equilibrate for 30 seconds during a “soak” step. Scaffolds were then stretched at a constant linear rate of 0.005 mm/s, and the slope of the initial linear portion of the stress-strain curve between 1-5% strain was calculated to measure stiffness. Tensile testing was performed for 300 seconds or until 25% strain if the scaffold yielded before 300 seconds.

### Myogenic Gene Expression Analysis via RT-qPCR

Mouse PCR primers for Pax3, Pax7, MyoD, Myf5, MyoG, and GAPDH (Integrated DNA Technology, IDT, Coralville, IA) are summarized in the supplemental information (**Table S1**).^57,58^ Myoblasts were differentiated for 5 days as described above. For RNA extraction, 500 µL of TRIzol™ Reagent (Fisher Scientific) was added to each well and incubated for 5 minutes at room temperature. The solution was collected in a microcentrifuge tube, and 200 µL of chloroform (Fisher Scientific) were added to each tube. The tube was vortexed and incubated on ice for 10 minutes before centrifuging at 12,000 rcf for 15 minutes at 4°C. After phase separation, the top aqueous phase was carefully collected into a new tube, and 250 µL of ethanol (Decon Labs, Prussia, PA) were added. The solution was transferred to a Zymo-spin column (Zymo Research, Irvine, CA) to collect the RNA. DNAse I treatment was performed according to the manufacturer’s instructions. RNA was then washed and eluted. The RNA quality was assessed using UV-vis spectroscopy at 260/280 nm and 260/230 nm (Implen NanoPhotometer). RNA was transcribed into cDNA using the LunaScript RT SuperMix Kit (New England Biolabs, Ipswich, MA), and controls without reverse transcriptase and the template were included to confirm successful cDNA transcription. Samples were stored at -20°C before proceeding to quantitative reverse transcription polymerase chain reaction (RT-qPCR). Quantabio PerfeCTa® SYBR® Green SuperMix, Low ROX™ (Quantabio, Beverly, MA) was used to perform RT-qPCR in a 10 µL reaction volume following the suggested reaction protocol using the QuantStudio5 Real-Time PCR System (ThermoFisher Scientific, Waltham, MA).

### Statistical Analysis

Data were analyzed and graphed using GraphPad Prism version 10.1.1 (Boston, MA).

Normality and equal variances were checked. Two-way analysis of variance (ANOVA) was used followed by the appropriate post-hoc test. P values less than 0.05 were considered statistically significant.

## Results

### Fabrication of HA-coated MEW Scaffolds

HA polymers were modified to enable hydrazone crosslinking as previously described (**Fig. 1**).^46^ 100 kDa HA was oxidized to expose aldehyde groups (HA-Ox). The number of monomers modified with aldehyde groups, which was called the percent degree of modification (%DOM), was determined to be 51.0±10.5% using ^1^H-NMR, and the percent yield was 66.0±35.3%. Simultaneously, an ion exchange was performed with 100 kDa HA to form an intermediate product HA-TBA. This intermediate product allowed for HA to become soluble in organic solvents such as DMSO for the modification with norbornene functional groups through Boc_2_O coupling for peptide conjugation (HA-Nor). The %DOM with norbornene functional groups was determined to be 24.0±2.3% using ^1^H-NMR, and the percent yield was determined to be 40.4±5.1%. HA-Nor was then modified with adipic acid dihydrazide (ADH) to form HA-Nor-ADH. The %DOM was determined to be 40.5±2.3% using ^1^H-NMR, and the percent yield was determined to be 65.9±9.5%. For hydrogels containing RGD, a photo-initiated thiol-ene click chemistry reaction was performed between the norbornene functional group on HA-Nor-ADH and the terminal cysteine of the RGD peptide. 55.4±29.3 nmol of RGD were conjugated per mg HA as determined using an Ellman’s assay that quantified the disappearance of free thiols. The reaction resulted in approximately 1.1 mM of RGD conjugated per 25 µL of HA-Nor-ADH-RGD, which was less than the desired 2.5 mM of RGD, but still within the range of RGD concentrations that has been previously shown to support cell attachment and survival in hydrogels.^43,59^

**Fig. 1.**
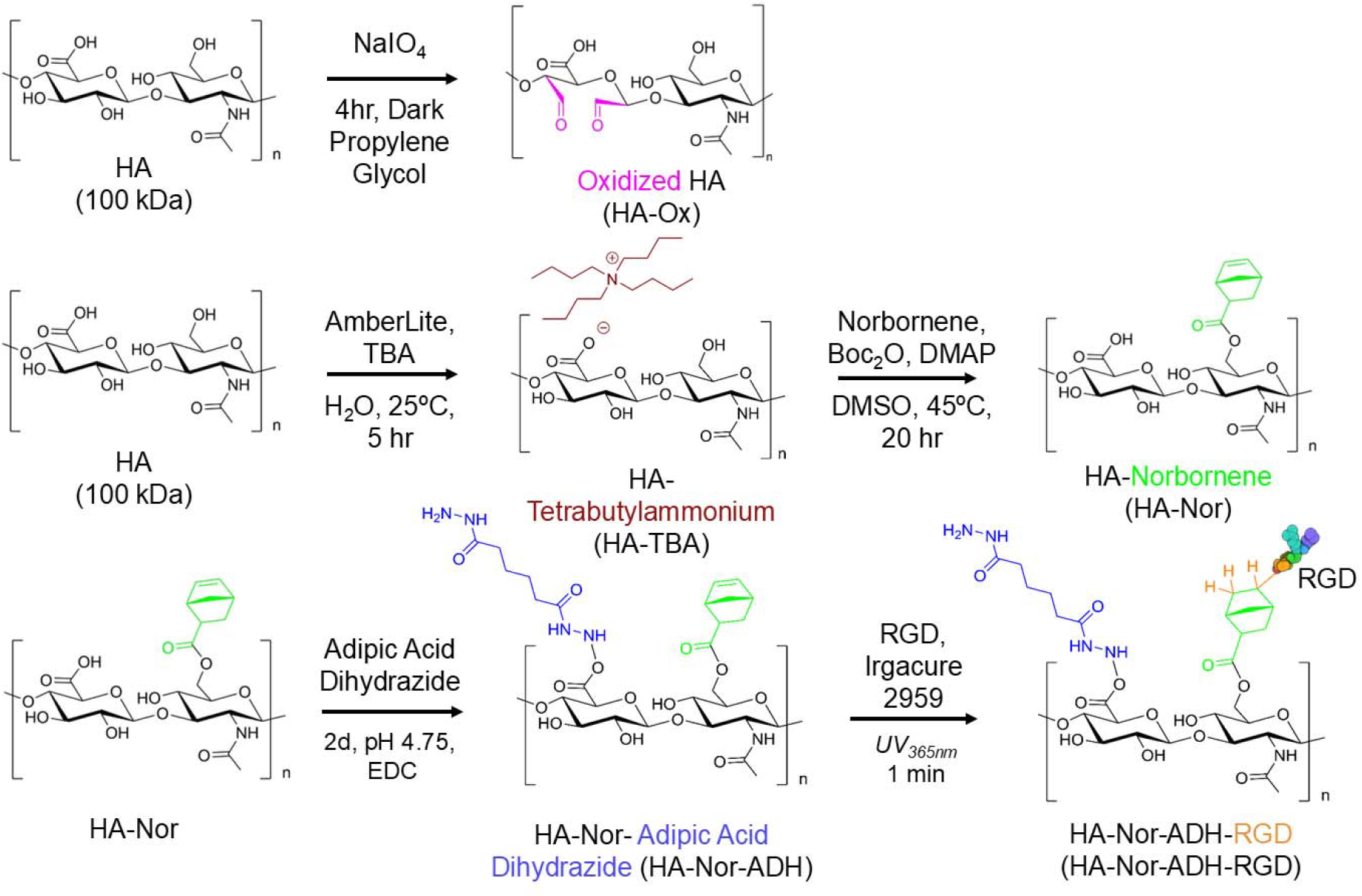
Hyaluronic acid (HA) modifications for hydrazone crosslinking. 100 kDa HA was either oxidized to form HA-Ox or modified with norbornene functional groups (HA-Nor). HA-Nor was then modified again with adipic acid dihydrazide functional groups to form HA-Nor-ADH. A thiol-ene click chemistry reaction was then performed with HA-Nor-ADH to form HA-Nor-ADH-RGD.

To coat scaffolds with the HA hydrogel, scaffolds were embedded between equal volumes of 2% w/v HA-Ox and 2% w/v HA-Nor-ADH or HA-Nor-ADH-RGD and allowed to gel for 30 minutes to create a thin hydrogel coating **(Fig. 2)**. The amine on ADH will spontaneously react with the aldehyde on HA-Ox through a Schiff base reaction to form hydrazone-crosslinked hydrogels.^46–48^

**Fig. 2.**
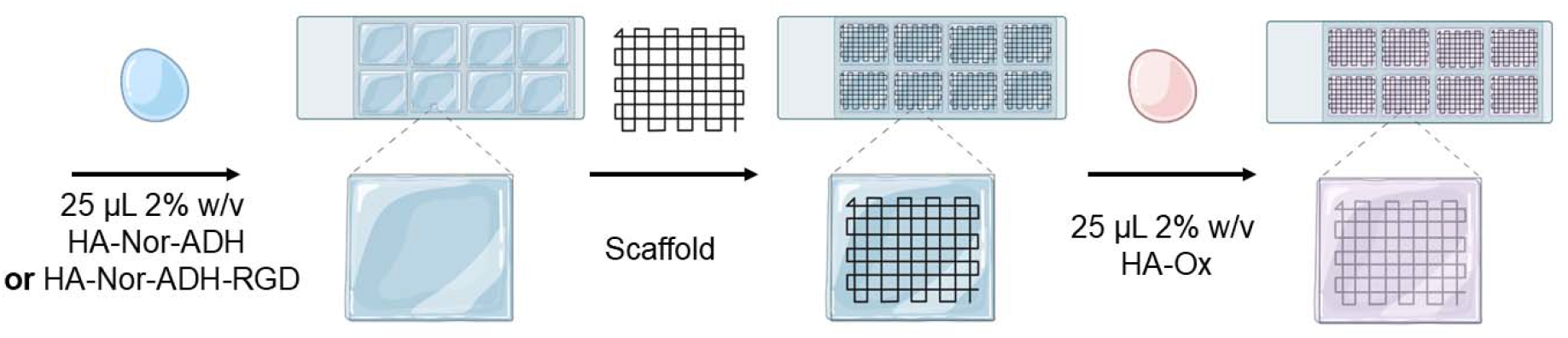
Coating of MEW scaffolds with HA. HA-coated scaffolds were fabricated in 8-chamber well plates by crosslinking 25 µL of either 2% w/v HA-Nor-ADH or HA-Nor-ADH-RGD and 25 µL of 2% w/v HA-Ox around MEW scaffolds for 30 minutes.

### Morphology of Non-coated and HA-coated MEW Scaffolds

MEW scaffolds were printed with either 10 µm, 20 µm, or 30 µm diameter PCL fibers to evaluate the effects of fiber diameter on myoblast attachment and differentiation. To create anisotropic cell alignment, myoblast alignment fibers were spaced 150 µm apart in the x-direction, and reinforcement fibers were spaced 500 µm apart in the y-direction to increase the number of fibers in a single direction while remaining within the printing limits of the machine. SEM images of non-coated MEW scaffolds revealed aligned fibers in both the x and y directions with varying fiber diameters (**Fig. 3A**). All fiber measurements are summarized in **Figure S2**. SEM images of lyophilized HA-coated scaffolds demonstrate coating of the HA hydrogel over the entire MEW scaffold, allowing fibers to remain visible (**Fig. 3B**).

**Fig. 3.**
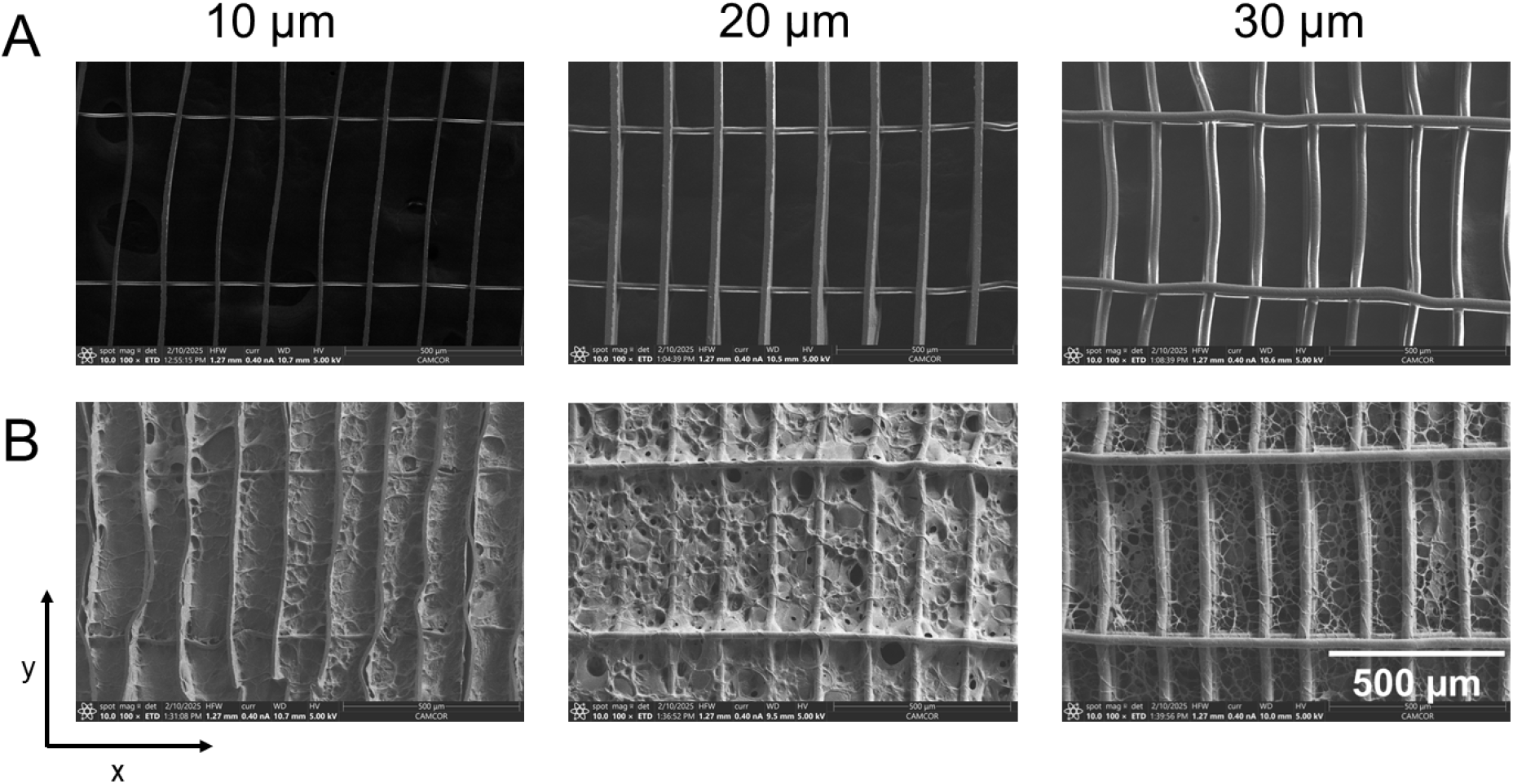
SEM images of non-coated and HA-coated MEW scaffolds. A) SEM images of non-coated MEW scaffolds with varying fiber diameters (10 µm, 20 µm, and 30 µm). B) SEM Images of HA-coated MEW scaffolds with varying fiber diameters (10 µm, 20 µm, and 30 µm). Scale bar = 500 µm. X and y directions as indicated for fiber measurements.

### Myoblast Attachment on MEW Scaffolds

C2C12 myoblasts were seeded on top of non-coated, HA-coated, or HA-RGD-coated MEW scaffolds to assess the impact of different coatings on cell attachment and alignment. Following 24 hours of proliferation, the cell culture medium was exchanged to low serum medium for 3 days to initiate the early phase of myogenic differentiation through serum starvation. Cells stained with Alexa Fluor™ 647 phalloidin for F-actin filaments and DAPI for cell nuclei revealed an increased presence of cells on the HA-coated and HA-RGD-coated scaffolds compared to the non-coated scaffolds (**Fig. 4**).

**Fig. 4.**
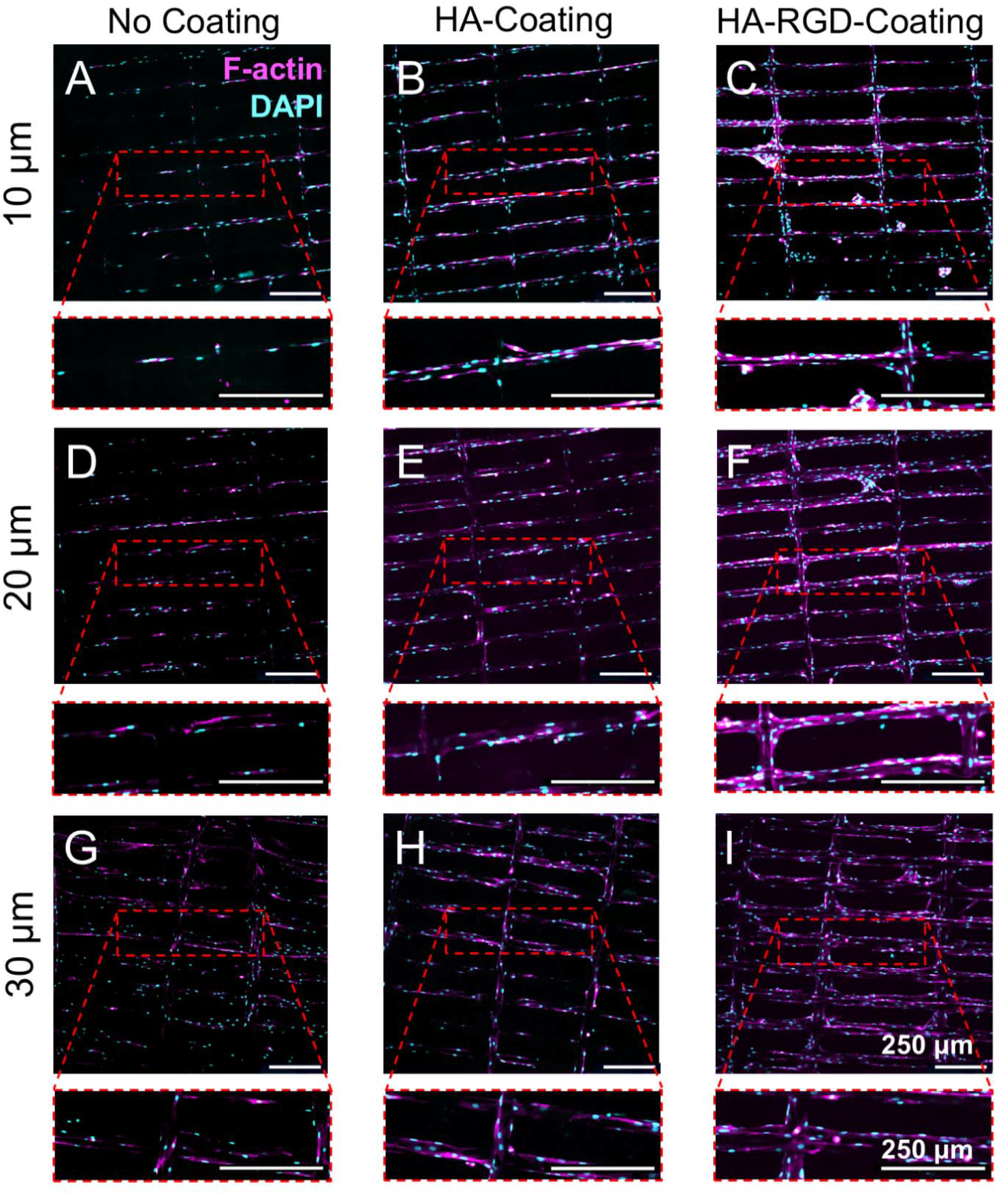
C2C12 myoblasts attach and align on MEW scaffolds. Immunocytochemistry images of C2C12 myoblasts on MEW scaffolds after four days of attachment. (pink = F-actin, blue = DAPI) Scale bars = 250 µm. A) 10 µm non-coated scaffolds, B) 10 µm HA-coated scaffolds, C) 10 µm HA-RGD-coated scaffolds, D) 20 µm non-coated scaffolds, E) 20 µm HA-coated scaffolds, F) 20 µm HA-RGD-coated scaffolds, G) 30 µm non-coated scaffolds, H) 30 µm HA-coated scaffolds, and I) 30 µm HA-RGD-coated scaffolds.

Cellular attachment was quantified by the number of nuclei, and cell alignment was determined by measuring nuclei directionality. The HA coating significantly increased cell attachment compared to non-coated scaffolds, and the HA-RGD coating further increased cell attachment compared to both HA-coated and non-coated scaffolds for all fiber diameters (**Fig. 5A**). Interestingly, there was also an effect of fiber diameter on myoblast attachment on HA-RGD-coated scaffolds with scaffolds containing 20 µm fibers having the highest nuclei count compared to scaffolds containing 10 µm and 30 µm fibers. Nuclei did not preferentially align in any direction on the non-coated and HA-coated scaffolds, as indicated by uniform nuclei directionality in all directions ranging from -90 to 90 degrees (**Fig. 5B-D**). Conversely, nuclei directionality was the highest at 0 degrees along fibers spaced 150 µm apart as intended for anisotropic myoblast alignment on HA-RGD-coated scaffolds containing 10 µm and 20 µm fibers (**Fig. 5B-C**).

**Fig. 5.**
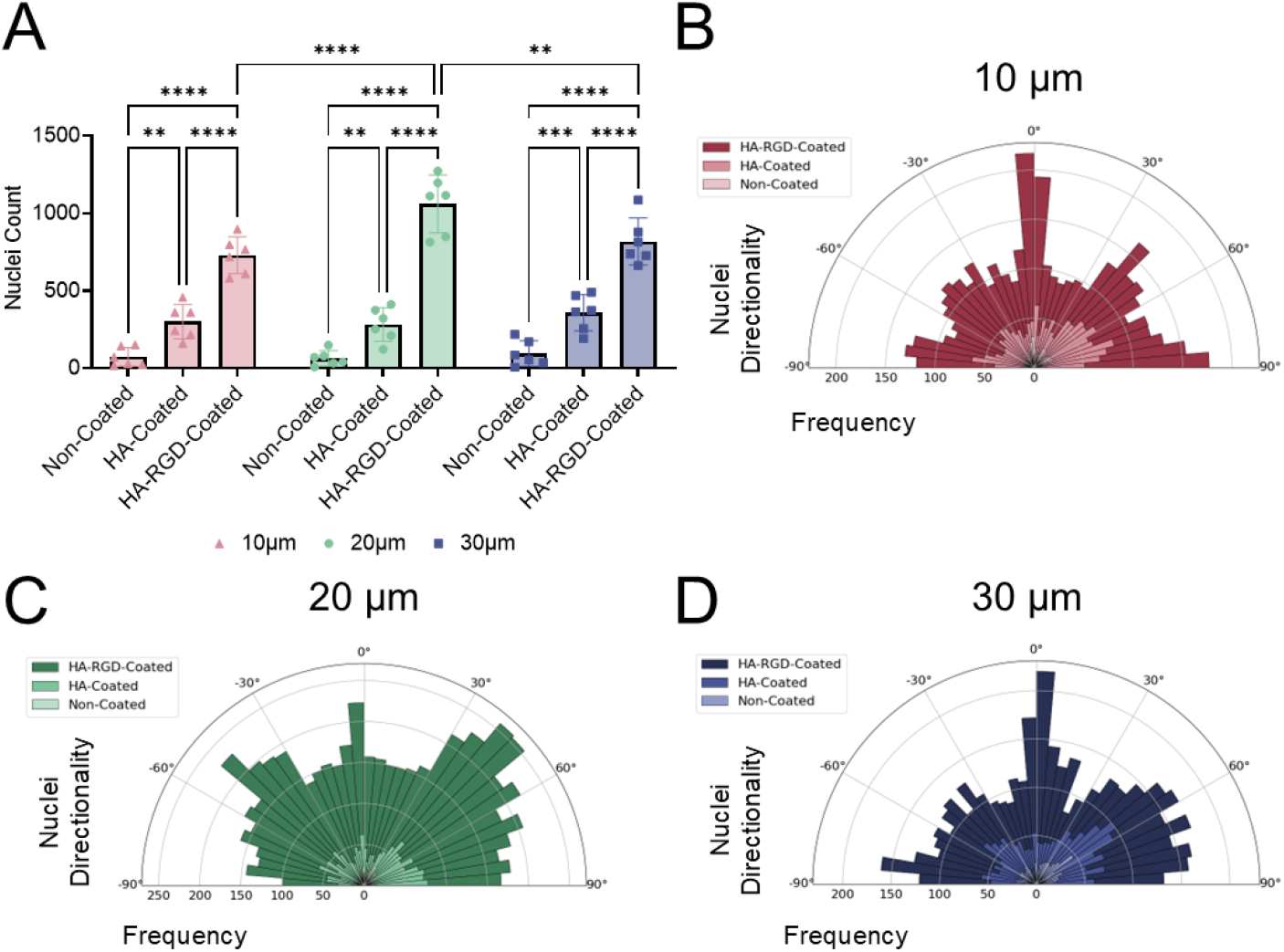
HA and HA-RGD-coatings increase myoblast attachment and anisotropic alignment on MEW scaffolds. A) Nuclei count on non-coated, HA-coated, and HA-RGD-coated scaffolds, n=6. (Two-way ANOVA with post-hoc Tukey’s multiple comparisons test, ^*^ p < 0.05, ^**^ p < 0.01, ^****^ p < 0.0001 as indicated) Nuclei directionality on non-coated, HA-coated, and HA-RGD-coated scaffolds with B) 10 µm, C) 20 µm, and D) 30 µm fibers, n=6. For 10 µm fiber scaffolds: non-coated 18-152 nuclei, HA-coated 162-458 nuclei, and HA-RGD-coated 583-898 nuclei. For 20 µm fiber scaffolds: non-coated 8-150 nuclei, HA-coated 122-412 nuclei, and HA-RGD-coated 815-1273 nuclei. For 30 µm fiber scaffolds: non-coated 8-219 nuclei, HA-coated 193-491 nuclei, and HA-RGD-coated 663-1087 nuclei.

### Myotube Formation on MEW Scaffolds

C2C12 myoblasts were differentiated on non-coated, HA-coated, or HA-RGD-coated MEW scaffolds to evaluate the effects of the various coatings on myotube formation. Following 24 hours of proliferation, the cell culture medium was exchanged to differentiation medium containing 2% v/v of horse serum for 7 days to promote mature myotube formation through serum starvation. Cells were then fixed and stained for myosin heavy chain (MHC), which is a late myogenic marker for myotubes, and DAPI to visualize the cell nuclei. Confocal imaging revealed an increase in cellular attachment and myotube formation on both HA-coated and HA-RGD-coated scaffolds (**Fig. 6**).

**Fig. 6.**
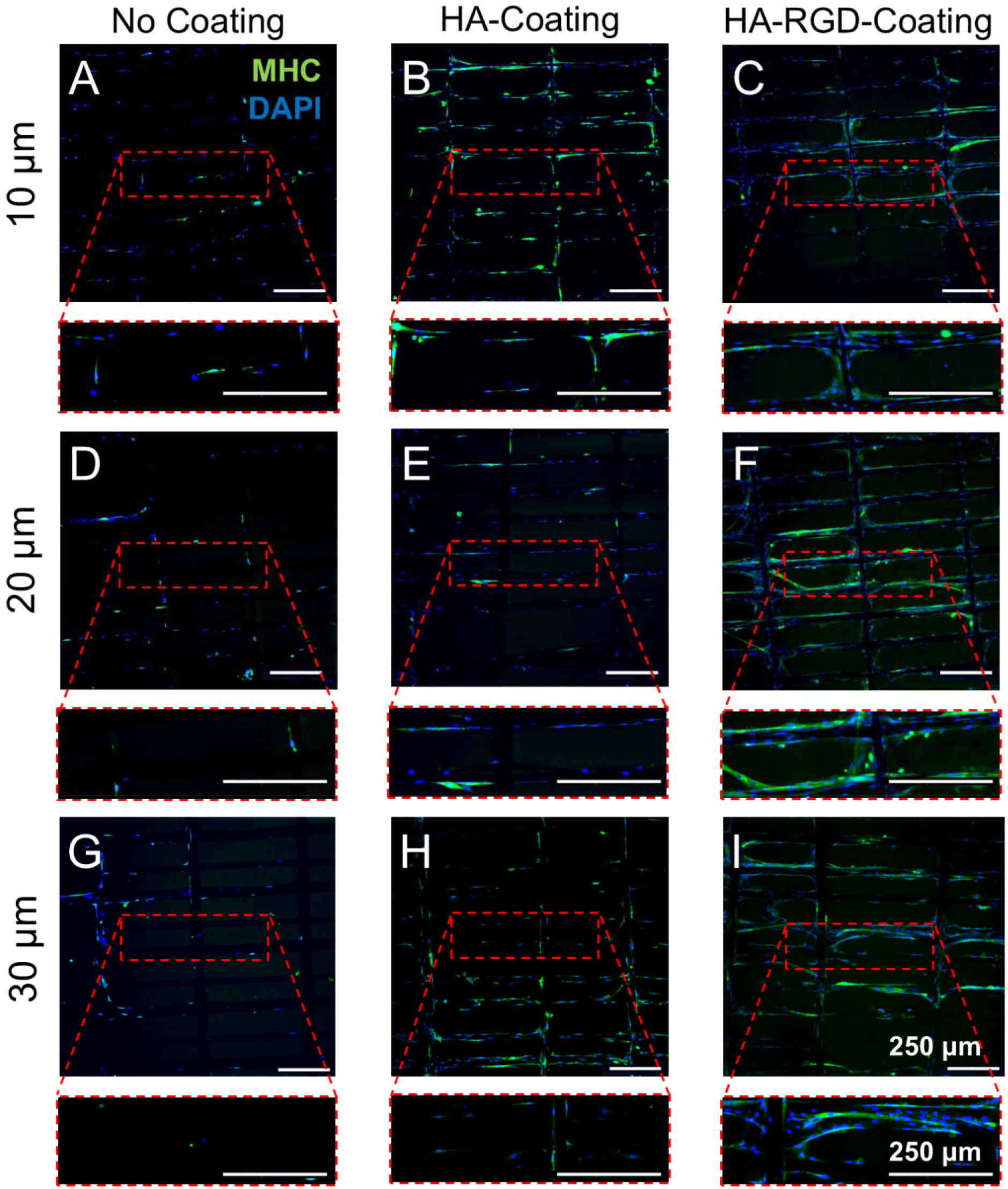
C2C12 myoblasts form aligned myotubes on MEW scaffolds. Immunocytochemistry images of C2C12 myoblasts differentiated on MEW scaffolds for 7 days, depicting myosin heavy chain (green) and DAPI (blue). Scale bar = 250 µm. A) 10 µm non-coated scaffolds, B) 10 µm HA-coated scaffolds, C) 10 µm HA-RGD-coated scaffolds, D) 20 µm non-coated scaffolds, E) 20 µm HA-coated scaffolds, F) 20 µm HA-RGD-coated scaffolds, G) 30 µm non-coated scaffolds, H) 30 µm HA-coated scaffolds, I) 30 µm HA-RGD-coated scaffolds.

Similar to results at early time points, nuclei counts revealed increased cell attachment on HA-coated scaffolds compared to non-coated scaffolds for all fiber diameters (**Fig. 7A**). The HA-RGD coating further increased cellular attachment compared to both non-coated and HA-coated scaffolds for all fiber diameters. HA-RGD-coated scaffolds containing 20 µm fibers demonstrated the highest cellular attachment compared to scaffolds containing 10 µm and 30 µm fibers. Cells cultured on both HA-coated and HA-RGD-coated 10 µm fiber scaffolds also demonstrated increased myotube length compared to non-coated 10 µm fiber scaffolds (**Fig. 7B**). However, no differences in myotube length were observed between scaffold coatings on 20 µm and 30 µm fiber scaffolds. The directionality of the myotubes in each scaffold was evaluated based on MHC staining. In all cases except non-coated 10 µm fiber scaffolds, increased myotube formation was observed in a single direction along horizontal fibers at 0 degrees regardless of the scaffold coating and fiber diameter (**Figure 7C-E**).

**Fig. 7.**
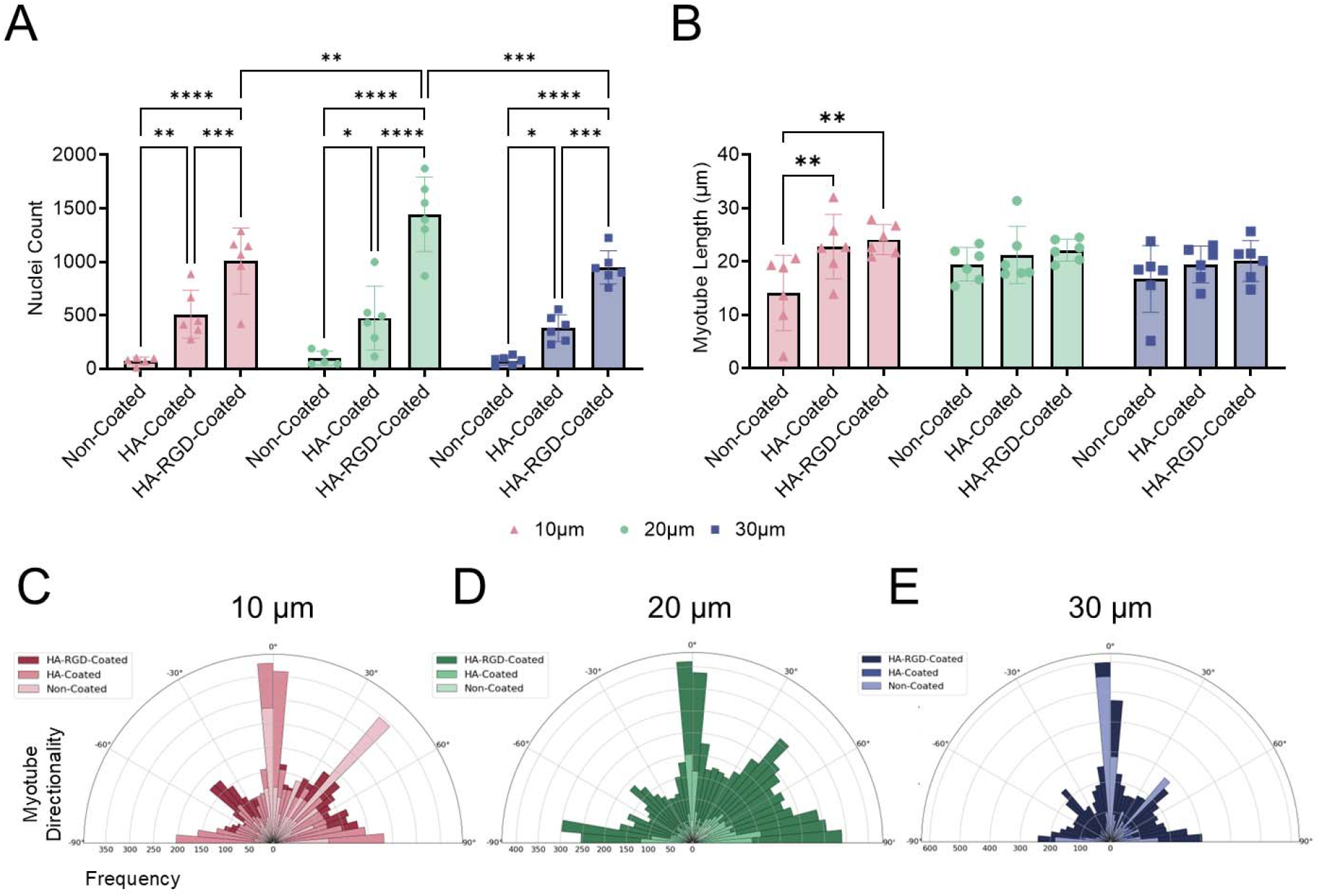
MEW scaffolds support anisotropic myotube formation, while HA and HA-RGD coatings increase cellular attachment. A) Nuclei count on non-coated, HA-coated, and HA-RGD-coated scaffolds containing 10 µm, 20 µm, and 30 µm diameter fibers, n=6. (Two-way ANOVA with post-hoc Tukey’s multiple comparisons test, ^*^ p < 0.05, ^**^ p < 0.01, ^***^ p < 0.001, ^****^ p < 0.0001 as indicated) B) Myotube length on non-coated, HA-coated, and HA-RGD-coated scaffolds containing 10 µm, 20 µm, and 30 µm diameter fibers, n=6. (Two-way ANOVA with post-hoc Tukey’s multiple comparisons test, ^**^ p < 0.01 as indicated) Myotube directionality for non-coated, HA-coated, and HA-RGD-coated scaffolds containing A) 10 µm, B) 20 µm, and C) 30 µm diameter fibers, n=6. For 10 µm fiber scaffolds: non-coated 50-2006 myotubes, HA-coated 301-1933 myotubes, and HA-RGD-coated 470-1386 myotubes. For 20 µm fiber scaffolds: non-coated 107-243 myotubes, HA-coated 194-670 myotubes, and HA-RGD-coated 658-2094 myotubes. For 30 µm fiber scaffolds: non-coated 102-2577 myotubes, HA-coated 179-596 myotubes, and HA-RGD-coated 464-4120 myotubes.

Since HA-RGD-coated scaffolds containing 20 µm fibers displayed higher cellular attachment than 10 and 30 µm scaffolds, we decided to move forward with these scaffolds for further analysis, including scaffold mechanical properties, creatine kinase activity, myogenic gene expression, and alpha-actinin staining to visualize sarcomeres.

### Mechanical Properties

Tensile testing was performed on non-coated MEW scaffolds containing 20 µm fibers for comparison to the mechanical properties of native skeletal muscle tissue. Non-coated scaffolds exhibited a stiffness of 0.843 ± 0.219 MPa (**Fig. S3**). In comparison, the stiffness of parallel human skeletal muscle fibers has been found to be approximately 0.44 MPa,^60^ suggesting that these scaffolds may provide similar mechanical support for myotube formation.

### Creatine Kinase Activity

Creatine kinase (CK) is an enzyme that facilitates the reversible transformation of creatine and ATP into phosphocreatine and adenosine diphosphate, which is important for muscle function and regeneration.^61^ When myoblasts start differentiating, they stop dividing and enter a final stage of development at which CK levels increase. As myoblasts fuse into myotubes, CK activity reaches its peak, making it a key marker of muscle cell differentiation and maturation.^62,63^ CK activity in cell-conditioned media was measured on day 1, day 5, and day 7 of differentiation to assess myogenic differentiation on non-coated, HA-coated, and HA-RGD-coated scaffolds, as well as cells cultured on tissue culture plastic (**Fig. 8**). No differences were observed in CK activity between scaffolds at individual time points. CK activity increased from day 1 to 5 for cells differentiated on HA-RGD-coated scaffolds and tissue culture plastic, indicating the formation of myotubes. CK activity then decreased from day 5 to 7 for cells differentiated on HA-coated scaffolds, HA-RGD-coated scaffolds, and tissue culture plastic. The subsequent decrease in CK levels may be due to the absence of growth factors that are needed to further promote the survival and maturation of myotubes.^64^ Overall, the highest levels of CK activity were observed on day 5. Along with the expression of MHC (**Fig. 6**), these findings further confirm the formation of myotubes on MEW scaffolds.

**Fig. 8.**
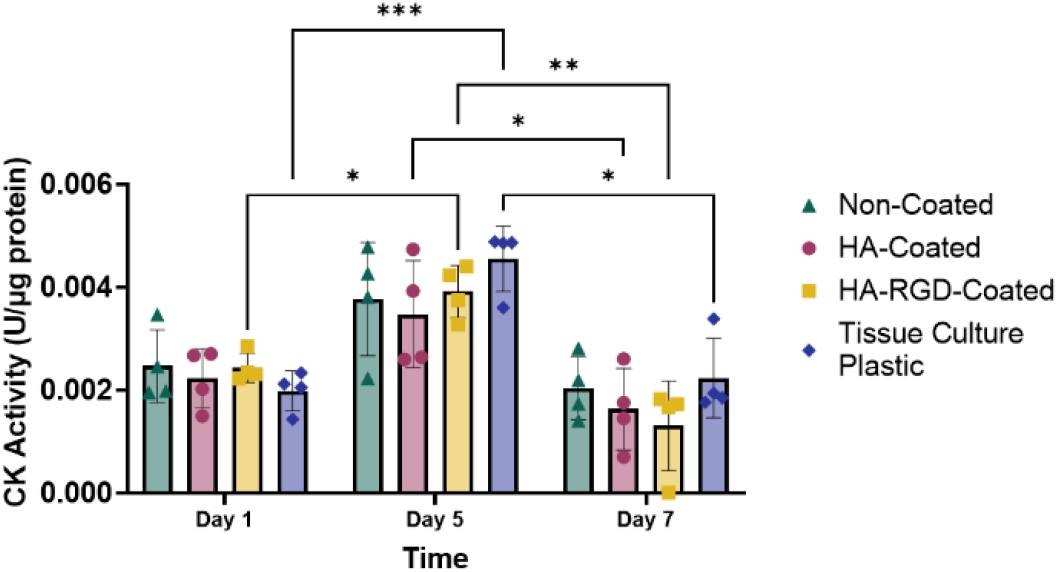
Creative kinase (CK) activity of myoblasts during differentiation. CK activity of C2C12 myoblasts differentiated on 20 µm fiber MEW scaffolds and tissue culture plastic after 1, 5, and 7 days of differentiation, n=4. (Two-way ANOVA post-hoc Tukey’s multiple comparisons test, ^*^ p < 0.05, ^**^ p < 0.01, ^***^ p < 0.001).

### Myogenic Gene Expression

Quantitative reverse transcription polymerase chain reaction (RT-qPCR) was performed on C2C12 myoblasts differentiated for 5 days on non-coated, HA-coated, and HA-RGD-coated 20 µm fiber scaffolds as well as tissue culture plastic and compared to undifferentiated myoblasts to determine the effects of substrate and coating on myogenic gene expression (**Fig. S4**). Expression of the myogenic genes Pax3, Pax7 MyoD, Myf5, and MyoG was evaluated and compared to GAPDH as the housekeeping gene.^65^ The expression of Pax3 suppresses myogenic differentiation^66^; as expected, Pax3 gene expression was decreased in all groups compared to undifferentiated cells (**Fig. S4A**). Pax7, MyoD, and Myf5 are myogenic genes that are expressed in the early stages of myoblast differentiation^67,68^ and were increased in all groups compared to undifferentiated cells (**Fig. S4B-D**). MyoD expression was significantly increased in cells grown on HA-coated scaffolds compared to HA-RGD-coated scaffolds, suggesting that HA-coated scaffolds may further support myotube formation. Interestingly, MyoG, which is typically expressed at later stages of myogenic differentiation,^69^ was also increased in all groups compared to undifferentiated myoblasts, with cells grown on HA-coated scaffolds expressing significantly more MyoG than cells grown on non-coated scaffolds, potentially indicating further progression of myogenic differentiation (**Fig. S4E**).

### Sarcomere Staining

Alpha-actinin staining was performed to visualize sarcomere striations after 7 days of differentiation to characterize the maturity of the myotubes (**Fig. 10**). While some striations were present in cells grown on the 20 µm HA-coated scaffold (**Fig. 10B**) and MHC-expressing cells were observed on all scaffolds, the lack of striations in cells grown on both the non-coated (**Fig. 10A**) and HA-RGD coated scaffolds (**Fig. 10C**) suggest that these myotubes have not fully matured.

**Fig. 10.**
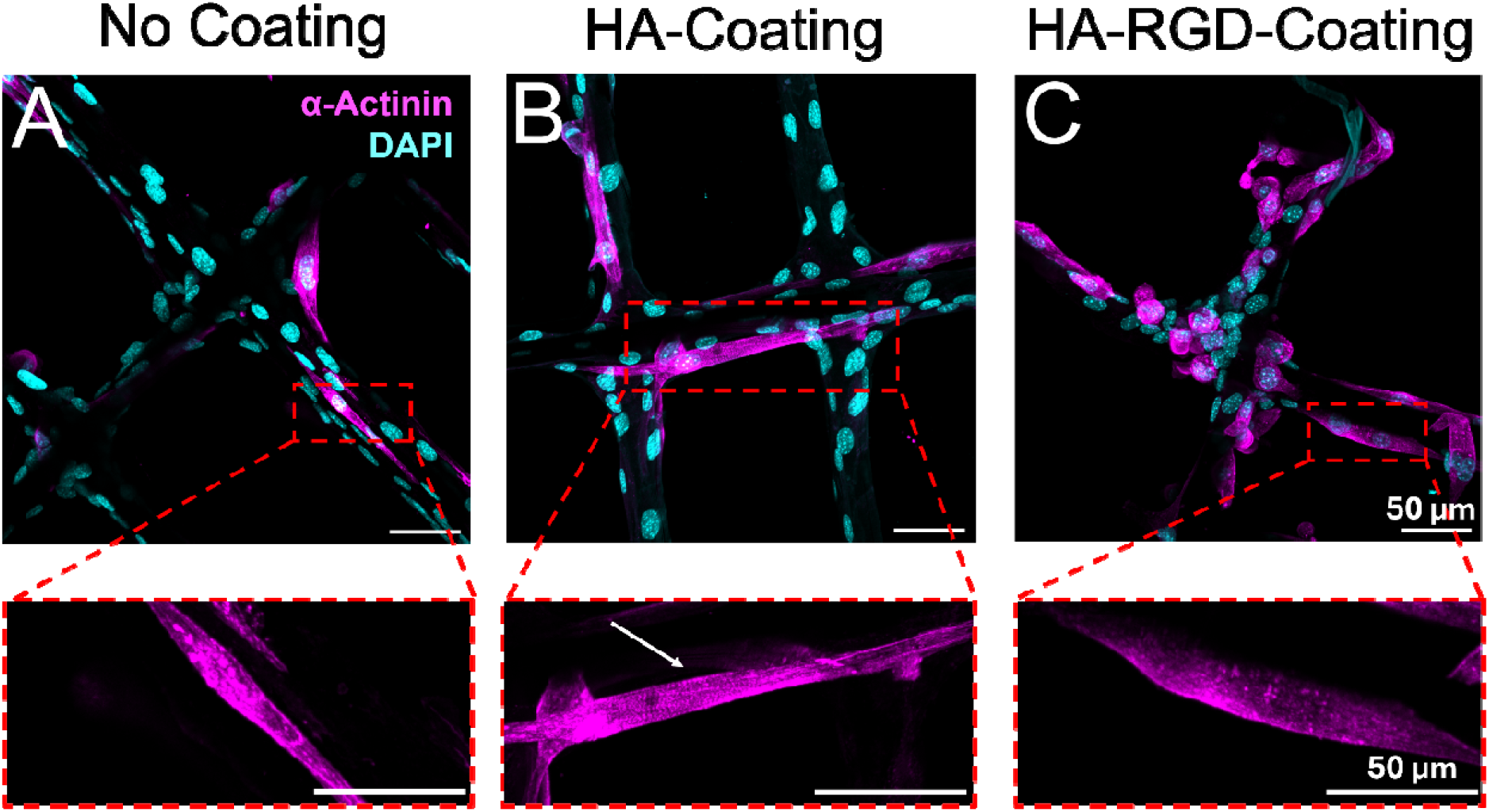
Alpha-actinin staining reveals presence of myotubes containing striations on HA-coated MEW scaffolds. Immunocytochemistry images of C2C12 myoblasts differentiated on 20 µm MEW scaffolds for 7 days, depicting alpha-actinin (pink) and DAPI (blue). Scale bar = 50 µm. A) non-coated scaffold, B) HA-coated scaffold with white arrow pointing to striations, C) HA-RGD-coated scaffold.

## Discussion

Composite materials containing both hydrogels and fiber-based scaffolds provide an opportunity to combine the advantages of the cell-supportive microenvironment of hydrogels with the alignment properties of fiber scaffolds. With their precisely aligned fibers, MEW scaffolds have great potential for directing cellular alignment for the repair of skeletal muscle tissue. Compared to electrospinning, a more commonly studied fiber-based printing technique in tissue engineering, MEW presents a finely tunable approach for exploring the effects of different fiber thickness and deposition geometries on anisotropic cell alignment.^45^ In this study, we harnessed the precision of MEW to fabricate scaffolds containing three different fiber diameters to study the effect of fiber diameter on myoblast alignment and myotube formation. We observed that 20 µm diameter fibers were preferred over 10 and 30 µm fiber diameters, which we hypothesize may be due to the similar size of the 20 µm diameter fibers to C2C12 myotubes, which range from 5 to 25 µm.^70,71^ MEW scaffolds made from PCL microfibers without proteins or cell-adhesive peptides have a limited ability to support myoblast attachment and differentiation themselves, revealing the shortcomings of synthetic polymer fiber-based scaffolds for muscle tissue engineering compared to scaffolds derived from natural ECM-based materials. For instance, Tan et al. demonstrated that the biophysical cues of an aligned type I collagen scaffold promoted a myogenic response that was dependent on the fibril size, porosity, and stiffness of the scaffold, further emphasizing the importance of considering ECM composition when directing cellular behavior with ECM-based scaffolds.^34^ However, while the cellular microenvironment provided by ECM-based scaffolds is amendable to myoblast differentiation, such scaffolds have a limited ability for tuning their physical properties without negatively impacting their biological activity and scaffold integrity. Moreover, it can be challenging to fabricate natural fiber scaffolds with the same precision provided by synthetic fiber scaffolds that can be fabricated using MEW.

We demonstrated that the cell microenvironment created by our MEW scaffolds could be significantly enhanced with the addition of a naturally-derived polymer matrix and ECM-derived cell-adhesive peptide. Our work highlights the value of combining biochemical cues with aligned fiber-based scaffolds to enhance material-cell interactions and provide a more supportive environment for cell growth. In previous work, we have demonstrated that our hydrazone-crosslinked HA hydrogel has a tunable gelation time, low swelling ratio, and stability over 28 days *in vitro*, making it an ideal hydrogel for creating a stable coating on MEW microfiber scaffolds.^46^ MEW scaffolds combined with HA and HA-RGD coatings exhibited increased myoblast attachment, myoblast alignment, and myotube formation compared to non-coated scaffolds, likely due to increased interactions between the cells, biopolymer, and cell-adhesive ligands.^49^

Our HA hydrogel has the additional advantage of being functionalized through orthogonal chemistry with norbornene functional groups for the bioconjugation of peptides and proteins. The effects of the HA coating on the MEW scaffolds were further enhanced with the addition of RGD, which increased myoblast attachment and alignment, and accelerated myotube formation. Our findings confirm results from previous studies, in which a similar concentration of RGD increased muscle satellite cell survival and differentiation in PEG-MAL hydrogels compared to the laminin-derived peptides YIGSR and C16.^43^ However, in other studies, the laminin-derived peptide IKVAV increased cell attachment and migration in PEG-DA-HA hydrogels compared to RGD.^20^ The presence of ECM peptides in HA hydrogels may also be important for myotube formation since the polyanionic surface of crosslinked HA hydrogels has been shown to decrease cell attachment.^20,72^ However, in our studies, we did not observe any limitations in cellular attachment on HA-coated scaffolds, which performed better in this regard than non-coated scaffolds. These positive cell attachment results may be due to the physical cues from the PCL fibers guiding cellular alignment and the viscoelastic properties of HA hydrazone-crosslinked hydrogels allowing for cell migration through the biomaterial. Regardless, HA-RGD-coated scaffolds significantly increased myoblast attachment, myoblast alignment, and myotube formation, although there were no differences on myotube length compared to HA-coated scaffolds.

Gene expression analysis revealed increased expression of early (Pax7, MyoD, Myf5) and late (MyoG) myogenic markers in myoblasts differentiated on all scaffolds and on tissue culture plastic, suggesting these cells were at an intermediate stage of differentiation. The presence of myosin heavy chain staining on all scaffolds further indicated robust myotube formation. However, we only observed some muscle striations on HA-coated scaffolds and not on the other scaffolds, suggesting that the myotubes on our scaffolds were not fully mature. We believe the lack of mature myotubes in our system may be due to the absence of exogenous growth factors, such as insulin-like growth factor 1 (IGF-1), basic fibroblast growth factor (bFGF), and epidermal growth factor (EGF), that may be necessary to promote further myogenic differentiation.^73–76^ The controlled release of growth factors, including IGF-1 and bFGF, has shown to promote muscle satellite cell proliferation and differentiation *in vitro* and improve volumetric muscle loss and fibrosis *in vivo* in a murine VML model.^77^

As MEW technology advances, we will continue to improve the scaffold design to optimize muscle healing outcomes. Minimizing the need for reinforcement fibers and increasing the number of fibers in a single direction will improve anisotropic myotube alignment, potentially enhancing muscle tissue repair. Furthermore, expanding on materials beyond PCL, such as PCL polypyrrole (PPy) or poly(3,4-ethylenedioxythiophene) (PEDOT) composite materials that have electroconductive properties, can further enhance muscle tissue repair, as it has been shown that electroconductive materials can activate cellular signaling related to proliferation and differentiation in electrically active tissues such as muscle.^78–80^ In addition, it would be motivating to investigate the effects of additional biochemical cues on myoblast attachment, alignment, and differentiation through the conjugation of other peptides and proteins to the HA backbone via the versatile norbornene functional group.^46,50,51^

Although there are limitations in the use of immortalized C2C12 myoblasts compared to primary myoblasts, including their ability to mimic the in vivo environment,^81^ we were able to use this cell line to investigate preliminary cellular behavior on HA-coated scaffolds, motivating future work to evaluate the effects of HA-coated scaffolds on primary muscle cell culture *in vitro*. In the future, we plan to evaluate HA-coated scaffolds as a muscle tissue repair patch *in vivo* in a rodent model of volumetric muscle loss in the quadriceps,^82,83^ which will provide further insight into the ability of this scaffold to repair muscle in a complex multicellular environment. Lastly, exploring the impact of the sustained delivery of growth factors, including IGF-1 and bFGF to promote myotube formation and maturation, may further enhance the utility of this scaffold in muscle tissue engineering applications.^46,77^

## Conclusion

Our HA-coated MEW microfiber scaffolds present a unique combination of biophysical and biochemical cues to enhance the attachment, alignment, and differentiation of skeletal myoblasts. The novelty of this work lies in the integration of MEW scaffolds with HA hydrogels, combining the benefits of both fiber-based scaffolds and natural ECM-based hydrogels. MEW scaffolds provide the structural framework for anisotropic cell alignment, while HA-RGD hydrogels offer the biochemical cues necessary for cell adhesion, alignment, and differentiation, allowing for enhanced cell-cell and cell-matrix interactions, which are essential for myogenic differentiation. While fully mature myotubes were not achieved with the use of our composite biomaterial, the main advantage of HA-coated and HA-RGD-coated MEW scaffolds lies in their ability to dramatically increase myoblast attachment and alignment, paving the way for future incorporation of growth factors and other cues to further enhance mature myotube formation and support eventual applications for functional muscle tissue formation.

## Acknowledgements

We are grateful for funding from the National Institutes of Health (R21 Trailblazer, R21-EB032112), Wu Tsai Human Performance Alliance, and Joe and Clara Tsai Foundation. J.D.K. was supported by the Knight Campus Undergraduate Scholars. P.D.D. was supported by the Bradshaw and Holzapfel Research Professor in Transformational Science and Mathematics. We are also grateful to Adam Fries for technical support on confocal imaging and analysis and Dr. David Johnson and Professor Danielle Benoit for technical support and use of the RSA-G2 Solids Analyzer.

## References

(1) Eugenis, I.; Wu, D.; Rando, T. A. Cells, Scaffolds, and Bioactive Factors: Engineering Strategies for Improving Regeneration Following Volumetric Muscle Loss. Biomaterials 2021, 278, 121173. 10.1016/j.biomaterials.2021.121173.

(2) Kim, J. T.; Kasukonis, B.; Brown, L.; Washington, T.; Wolchok, J. C. Recovery from Volumetric Muscle Loss Injury: A Comparison Between Young and Aged Rats. Exp Gerontol 2016, 83, 37–46. 10.1016/j.exger.2016.07.008.

(3) Volpi, M.; Paradiso, A.; Costantini, M.; Świ□szkowski, W. Hydrogel-Based Fiber Biofabrication Techniques for Skeletal Muscle Tissue Engineering. ACS Biomater Sci Eng 2022, 8 (2), 379–405. 10.1021/acsbiomaterials.1c01145.

(4) Gilbert-Honick, J.; Grayson, W. Vascularized and Innervated Skeletal Muscle Tissue Engineering. Adv Healthc Mater 2020, 9 (1), e1900626. 10.1002/adhm.201900626.

(5) Kim, J. T.; Kasukonis, B.; Brown, L.; Washington, T.; Wolchok, J. C. Recovery from Volumetric Muscle Loss Injury: A Comparison Between Young and Aged Rats. Exp Gerontol 2016, 83, 37–46. 10.1016/j.exger.2016.07.008.

(6) Purslow, P. P. The Structure and Role of Intramuscular Connective Tissue in Muscle Function. Front. Physiol. 2020, 11. 10.3389/fphys.2020.00495.

(7) Zhang, W.; Liu, Y.; Zhang, H. Extracellular Matrix: An Important Regulator of Cell Functions and Skeletal Muscle Development. Cell & Bioscience 2021, 11 (1), 65. 10.1186/s13578-021-00579-4.

(8) Ward, C. L.; Ji, L.; Corona, B. T. An Autologous Muscle Tissue Expansion Approach for the Treatment of Volumetric Muscle Loss. Biores Open Access 2015, 4 (1), 198–208. 10.1089/biores.2015.0009.

(9) Greising, S. M.; Warren, G. L.; Southern, W. M.; Nichenko, A. S.; Qualls, A. E.; Corona, B. T.; Call, J. A. Early Rehabilitation for Volumetric Muscle Loss Injury Augments Endogenous Regenerative Aspects of Muscle Strength and Oxidative Capacity. BMC Musculoskelet Disord 2018, 19, 173. 10.1186/s12891-018-2095-6.

(10) Mintz, E. L.; Passipieri, J. A.; Franklin, I. R.; Toscano, V. M.; Afferton, E. C.; Sharma, P. R.; Christ, G. J. Long-Term Evaluation of Functional Outcomes Following Rat Volumetric Muscle Loss Injury and Repair. Tissue Eng Part A 2020, 26 (3–4), 140–156. 10.1089/ten.TEA.2019.0126.

(11) Dolan, C. P.; Motherwell, J. M.; Franco, S. R.; Janakiram, N. B.; Valerio, M. S.; Goldman, S. M.; Dearth, C. L. Evaluating the Potential Use of Functional Fibrosis to Facilitate Improved Outcomes Following Volumetric Muscle Loss Injury. Acta Biomaterialia 2022, 140, 379–388. 10.1016/j.actbio.2021.11.032.

(12) Hu, C.; Chiang, G.; Chan, A. H.-P.; Alcazar, C.; Nakayama, K. H.; Quarta, M.; Rando, T. A.; Huang, N. F. A Mouse Model of Volumetric Muscle Loss and Therapeutic Scaffold Implantation. Nat Protoc 2024. 10.1038/s41596-024-01059-y.

(13) Carnes, M. E.; Pins, G. D. Skeletal Muscle Tissue Engineering: Biomaterials-Based Strategies for the Treatment of Volumetric Muscle Loss. Bioengineering 2020, 7 (3), 85. 10.3390/bioengineering7030085.

(14) Niknezhad, S. V.; Mehrali, M.; Khorasgani, F. R.; Heidari, R.; Kadumudi, F. B.; Golafshan, N.; Castilho, M.; Pennisi, C. P.; Hasany, M.; Jahanshahi, M.; Mehrali, M.; Ghasemi, Y.; Azarpira, N.; Andresen, T. L.; Dolatshahi-Pirouz, A. Enhancing Volumetric Muscle Loss (VML) Recovery in a Rat Model Using Super Durable Hydrogels Derived from Bacteria. Bioactive Materials 2024, 38, 540–558. 10.1016/j.bioactmat.2024.04.006.

(15) Browe, D.; Freeman, J. Optimizing C2C12 Myoblast Differentiation Using Polycaprolactone-Polypyrrole Copolymer Scaffolds. J Biomed Mater Res A 2019, 107 (1), 220–231. 10.1002/jbm.a.36556.

(16) Carnes, M. E.; Pins, G. D. Skeletal Muscle Tissue Engineering: Biomaterials-Based Strategies for the Treatment of Volumetric Muscle Loss. Bioengineering (Basel) 2020, 7 (3), 85. 10.3390/bioengineering7030085.

(17) Zhu, C.; Sklyar, K.; Karvar, M.; Endo, Y.; Sinha, I. Scaffold Tissue Engineering Strategies for Volumetric Muscle Loss. par 2023, 10 (0), N/A-N/A. 10.20517/2347-9264.2022.89.

(18) Bian, W.; Juhas, M.; Pfeiler, T. W.; Bursac, N. Local Tissue Geometry Determines Contractile Force Generation of Engineered Muscle Networks. Tissue Eng Part A 2012, 18 (9–10), 957–967. 10.1089/ten.TEA.2011.0313.

(19) Narayanan, N.; Jia, Z.; Kim, K. H.; Kuang, L.; Lengemann, P.; Shafer, G.; Bernal-Crespo, V.; Kuang, S.; Deng, M. Biomimetic Glycosaminoglycan-Based Scaffolds Improve Skeletal Muscle Regeneration in a Murine Volumetric Muscle Loss Model. Bioactive Materials 2021, 6 (4), 1201–1213. 10.1016/j.bioactmat.2020.10.012.

(20) Silva Garcia, J. M.; Panitch, A.; Calve, S. Functionalization of Hyaluronic Acid Hydrogels with ECM-Derived Peptides to Control Myoblast Behavior. Acta Biomater 2019, 84, 169–179. 10.1016/j.actbio.2018.11.030.

(21) Hwangbo, H.; Lee, H.; Jin, E.-J.; Lee, J.; Jo, Y.; Ryu, D.; Kim, G. Bio-Printing of Aligned GelMa-Based Cell-Laden Structure for Muscle Tissue Regeneration. Bioactive Materials 2022, 8, 57–70. 10.1016/j.bioactmat.2021.06.031.

(22) Wang, L.; Wu, Y.; Guo, B.; Ma, P. X. Nanofiber Yarn/Hydrogel Core–Shell Scaffolds Mimicking Native Skeletal Muscle Tissue for Guiding 3D Myoblast Alignment, Elongation, and Differentiation. ACS Nano 2015, 9 (9), 9167–9179. 10.1021/acsnano.5b03644.

(23) Abdalkader, R.; Konishi, S.; Fujita, T. The Development of Biomimetic Aligned Skeletal Muscles in a Fully 3D Printed Microfluidic Device. Biomimetics (Basel) 2021, 7 (1), 2. 10.3390/biomimetics7010002.

(24) Hu, T.; Shi, M.; Zhao, X.; Liang, Y.; Bi, L.; Zhang, Z.; Liu, S.; Chen, B.; Duan, X.; Guo, B. Biomimetic 3D Aligned Conductive Tubular Cryogel Scaffolds with Mechanical Anisotropy for 3D Cell Alignment, Differentiation and in Vivo Skeletal Muscle Regeneration. Chemical Engineering Journal 2022, 428, 131017. 10.1016/j.cej.2021.131017.

(25) Cheng, Y.-W.; Shiwarski, D. J.; Ball, R. L.; Whitehead, K. A.; Feinberg, A. W. Engineering Aligned Skeletal Muscle Tissue Using Decellularized Plant-Derived Scaffolds. ACS Biomater Sci Eng 2020, 6 (5), 3046–3054. 10.1021/acsbiomaterials.0c00058.

(26) Chen, H.; Zhong, J.; Wang, J.; Huang, R.; Qiao, X.; Wang, H.; Tan, Z. Enhanced Growth and Differentiation of Myoblast Cells Grown on E-Jet 3D Printed Platforms. Int J Nanomedicine 2019, 14, 937–950. 10.2147/IJN.S193624.

(27) Pacilio, S.; Costa, R.; Papa, V.; Rodia, M. T.; Gotti, C.; Pagnotta, G.; Cenacchi, G.; Focarete, M. L. Electrospun Poly(L-Lactide-Co-ε-Caprolactone) Scaffold Potentiates C2C12 Myoblast Bioactivity and Acts as a Stimulus for Cell Commitment in Skeletal Muscle Myogenesis. Bioengineering (Basel) 2023, 10 (2), 239. 10.3390/bioengineering10020239.

(28) Nakayama, K. H.; Shayan, M.; Huang, N. F. Engineering Biomimetic Materials for Skeletal Muscle Repair and Regeneration. Adv Healthc Mater 2019, 8 (5), e1801168. 10.1002/adhm.201801168.

(29) Zhang, S.; Guo, Y.; Lu, Y.; Liu, F.; Heng, B. C.; Deng, X. The Considerations on Selecting the Appropriate Decellularized ECM for Specific Regeneration Demands. Mater Today Bio 2024, 29, 101301. 10.1016/j.mtbio.2024.101301.

(30) Chaturvedi, V.; Dye, D. E.; Kinnear, B. F.; van Kuppevelt, T. H.; Grounds, M. D.; Coombe, D. R. Interactions between Skeletal Muscle Myoblasts and Their Extracellular Matrix Revealed by a Serum Free Culture System. PLoS One 2015, 10 (6), e0127675. 10.1371/journal.pone.0127675.

(31) Kozan, N. G.; Joshi, M.; Sicherer, S. T.; Grasman, J. M. Porous Biomaterial Scaffolds for Skeletal Muscle Tissue Engineering. Front Bioeng Biotechnol 2023, 11, 1245897. 10.3389/fbioe.2023.1245897.

(32) Colin, C.; Akpo, E.; Perrin, A.; Cornu, D.; Cambedouzou, J. Encapsulation in Alginates Hydrogels and Controlled Release: An Overview. Molecules 2024, 29 (11), 2515. 10.3390/molecules29112515.

(33) Kozan, N. G.; Caswell, S.; Patel, M.; Grasman, J. M. Aligned Collagen Sponges with Tunable Pore Size for Skeletal Muscle Tissue Regeneration. J Funct Biomater 2023, 14 (11), 533. 10.3390/jfb14110533.

(34) Tan, Y. H.; Habing, K. M.; Riesterer, J. L.; Stempinski, E. S.; Lewis, S. H.; Pfeifer, C. S.; Malhotra, S. V.; Nakayama, K. H. Engineered Nanofibrillar Collagen with Tunable Biophysical Properties for Myogenic, Endothelial, and Osteogenic Cell Guidance. Acta Biomater 2024, 186, 95–107. 10.1016/j.actbio.2024.08.002.

(35) Yang, G. H.; Kim, W.; Kim, J.; Kim, G. A Skeleton Muscle Model Using GelMA-Based Cell-Aligned Bioink Processed with an Electric-Field Assisted 3D/4D Bioprinting. Theranostics 2021, 11 (1), 48–63. 10.7150/thno.50794.

(36) Borselli, C.; Cezar, C. A.; Shvartsman, D.; Vandenburgh, H. H.; Mooney, D. J. The Role of Multifunctional Delivery Scaffold in the Ability of Cultured Myoblasts to Promote Muscle Regeneration. Biomaterials 2011, 32 (34), 8905–8914. 10.1016/j.biomaterials.2011.08.019.

(37) Panayi, A. C.; Smit, L.; Hays, N.; Udeh, K.; Endo, Y.; Li, B.; Sakthivel, D.; Tamayol, A.; Neppl, R. L.; Orgill, D. P.; Nuutila, K.; Sinha, I. A Porous Collagen-GAG Scaffold Promotes Muscle Regeneration Following Volumetric Muscle Loss Injury. Wound Repair Regen 2020, 28 (1), 61–74. 10.1111/wrr.12768.

(38) Hu, C.; Ayan, B.; Chiang, G.; Chan, A. H. P.; Rando, T. A.; Huang, N. F. Comparative Effects of Basic Fibroblast Growth Factor Delivery or Voluntary Exercise on Muscle Regeneration after Volumetric Muscle Loss. Bioengineering 2022, 9 (1), 37. 10.3390/bioengineering9010037.

(39) Saveh-Shemshaki, N.; Barajaa, M. A.; Otsuka, T.; Mirdamadi, E. S.; Nair, L. S.; Laurencin, C. T. Electroconductivity, a Regenerative Engineering Approach to Reverse Rotator Cuff Muscle Degeneration. Regen Biomater 2023, 10, rbad099. 10.1093/rb/rbad099.

(40) Wang, L.; Cao, L.; Shansky, J.; Wang, Z.; Mooney, D.; Vandenburgh, H. Minimally Invasive Approach to the Repair of Injured Skeletal Muscle with a Shape-Memory Scaffold. Mol Ther 2014, 22 (8), 1441–1449. 10.1038/mt.2014.78.

(41) Csapo, R.; Gumpenberger, M.; Wessner, B. Skeletal Muscle Extracellular Matrix – What Do We Know About Its Composition, Regulation, and Physiological Roles? A Narrative Review. Front Physiol 2020, 11, 253. 10.3389/fphys.2020.00253.

(42) Ibáñez-Fonseca, A.; Santiago Maniega, S.; Gorbenko del Blanco, D.; Catalán Bernardos, B.; Vega Castrillo, A.; Álvarez Barcia, Á. J.; Alonso, M.; Aguado, H. J.; Rodríguez-Cabello, J. C. Elastin-Like Recombinamer Hydrogels for Improved Skeletal Muscle Healing Through Modulation of Macrophage Polarization. Front Bioeng Biotechnol 2020, 8, 413. 10.3389/fbioe.2020.00413.

(43) Han, W. M.; Anderson, S. E.; Mohiuddin, M.; Barros, D.; Nakhai, S. A.; Shin, E.; Amaral, I. F.; Pêgo, A. P.; García, A. J.; Jang, Y. C. Synthetic Matrix Enhances Transplanted Satellite Cell Engraftment in Dystrophic and Aged Skeletal Muscle with Comorbid Trauma. Sci Adv 2018, 4 (8), eaar4008. 10.1126/sciadv.aar4008.

(44) Brown, T. D.; Dalton, P. D.; Hutmacher, D. W. Direct Writing by Way of Melt Electrospinning. Adv Mater 2011, 23 (47), 5651–5657. 10.1002/adma.201103482.

(45) Calero-Castro, F. J.; Perez-Puyana, V. M.; Laga, I.; Padillo Ruiz, J.; Romero, A.; de la Portilla de Juan, F. Mechanical Stimulation and Aligned Poly(ε-Caprolactone)-Gelatin Electrospun Scaffolds Promote Skeletal Muscle Regeneration. ACS Appl Bio Mater 2024, 7 (10), 6430–6440. 10.1021/acsabm.4c00559.

(46) Mozipo, E. A.; Galindo, A. N.; Khachatourian, J. D.; Harris, C. G.; Dorogin, J.; Spaulding, V. R.; Ford, M. R.; Singhal, M.; Fogg, K. C.; Hettiaratchi, M. H. Statistical Optimization of Hydrazone-Crosslinked Hyaluronic Acid Hydrogels for Protein Delivery. J. Mater. Chem. B 2024, 12 (10), 2523–2536. 10.1039/D3TB01588B.

(47) França, C. G.; Villalva, D. G.; Santana, M. H. A. Oxi-HA/ADH Hydrogels: A Novel Approach in Tissue Engineering and Regenerative Medicine. Polysaccharides 2021, 2 (2), 477–496. 10.3390/polysaccharides2020029.

(48) Su, W.-Y.; Chen, K.-H.; Chen, Y.-C.; Lee, Y.-H.; Tseng, C.-L.; Lin, F.-H. An Injectable Oxidated Hyaluronic Acid/Adipic Acid Dihydrazide Hydrogel as a Vitreous Substitute. Journal of Biomaterials Science -- Polymer Edition 2011, 22 (13), 1777–1797. 10.1163/092050610X522729.

(49) Leng, Y.; Abdullah, A.; Wendt, M. K.; Calve, S. Hyaluronic Acid, CD44 and RHAMM Regulate Myoblast Behavior during Embryogenesis. Matrix Biol 2019, 78–79, 236–254. 10.1016/j.matbio.2018.08.008.

(50) Gramlich, W. M.; Kim, I. L.; Burdick, J. A. Synthesis and Orthogonal Photopatterning of Hyaluronic Acid Hydrogels with Thiol-Norbornene Chemistry. Biomaterials 2013, 34 (38), 9803–9811. 10.1016/j.biomaterials.2013.08.089.

(51) Davidson, M. D.; Ban, E.; Schoonen, A. C. M.; Lee, M.-H.; D’Este, M.; Shenoy, V. B.; Burdick, J. A. Mechanochemical Adhesion and Plasticity in Multifiber Hydrogel Networks. Advanced Materials 2020, 32 (8), 1905719. 10.1002/adma.201905719.

(52) Zhao, H.; Heindel, N. D. Determination of Degree of Substitution of Formyl Groups in Polyaldehyde Dextran by the Hydroxylamine Hydrochloride Method. Pharm Res 1991, 8 (3), 400–402. 10.1023/A:1015866104055.

(53) Wang, Q.; Wang, Z.; Zhang, D.; Gu, J.; Ma, Y.; Zhang, Y.; Chen, J. Circular Patterns of Dynamic Covalent Hydrogels with Gradient Stiffness for Screening of the Stem Cell Microenvironment. ACS Appl. Mater. Interfaces 2022, 14 (42), 47461–47471. 10.1021/acsami.2c14924.

(54) Dalton, P. D.; Polites, M. Visualizing fiber path and generating G-code for melt electrowriting of tubular scaffolds using Grasshopper software.

(55) Hrynevich, A.; Achenbach, P.; Jungst, T.; Brook, G. A.; Dalton, P. D. Design of Suspended Melt Electrowritten Fiber Arrays for Schwann Cell Migration and Neurite Outgrowth. Macromol Biosci 2021, 21 (7), e2000439. 10.1002/mabi.202000439.

(56) Hettiaratchi, M. H.; Miller, T.; Temenoff, J. S.; Guldberg, R. E.; McDevitt, T. C. Heparin Microparticle Effects on Presentation and Bioactivity of Bone Morphogenetic Protein-2. Biomaterials 2014, 35 (25), 7228–7238. 10.1016/j.biomaterials.2014.05.011.

(57) Wang, H.; Lööf, S.; Borg, P.; Nader, G. A.; Blau, H. M.; Simon, A. Turning Terminally Differentiated Skeletal Muscle Cells into Regenerative Progenitors. Nat Commun 2015, 6 (1), 7916. 10.1038/ncomms8916.

(58) Investigation of niclosamide as a repurposing agent for skeletal muscle atrophy - PubMed. https://pubmed.ncbi.nlm.nih.gov/34038481/ (accessed 2025-07-28).

(59) Burdick, J. A.; Anseth, K. S. Photoencapsulation of Osteoblasts in Injectable RGD-Modified PEG Hydrogels for Bone Tissue Engineering. Biomaterials 2002, 23 (22), 4315–4323. 10.1016/S0142-9612(02)00176-X.

(60) Kuthe, C. D.; Uddanwadiker, R. V.; Ramteke, A. Experimental Evaluation of Fiber Orientation Based Material Properties of Skeletal Muscle in Tension. Mol Cell Biomech 2014, 11 (2), 113–128.

(61) Tai, P. W.; Fisher-Aylor, K. I.; Himeda, C. L.; Smith, C. L.; MacKenzie, A. P.; Helterline, D. L.; Angello, J. C.; Welikson, R. E.; Wold, B. J.; Hauschka, S. D. Differentiation and Fiber Type-Specific Activity of a Muscle Creatine Kinase Intronic Enhancer. Skeletal Muscle 2011, 1 (1), 25. 10.1186/2044-5040-1-25.

(62) Diel, P.; Baadners, D.; Schlüpmann, K.; Velders, M.; Schwarz, J. P. C2C12 Myoblastoma Cell Differentiation and Proliferation Is Stimulated by Androgens and Associated with a Modulation of Myostatin and Pax7 Expression. J Mol Endocrinol 2008, 40 (5), 231–241. 10.1677/JME-07-0175.

(63) Jang, M.; Scheffold, J.; Røst, L. M.; Cheon, H.; Bruheim, P. Serum-Free Cultures of C2C12 Cells Show Different Muscle Phenotypes Which Can Be Estimated by Metabolic Profiling. Sci Rep 2022, 12 (1), 827. 10.1038/s41598-022-04804-z.

(64) Cai, A.; Hardt, M.; Schneider, P.; Schmid, R.; Lange, C.; Dippold, D.; Schubert, D. W.; Boos, A. M.; Weigand, A.; Arkudas, A.; Horch, R. E.; Beier, J. P. Myogenic Differentiation of Primary Myoblasts and Mesenchymal Stromal Cells under Serum-Free Conditions on PCL-Collagen I-Nanoscaffolds. BMC Biotechnol 2018, 18 (1), 75. 10.1186/s12896-018-0482-6.

(65) Ma, J.; Chen, J.; Gan, M.; Chen, L.; Zhao, Y.; Niu, L.; Zhu, Y.; Zhang, S.; Li, X.; Guo, Z.; Wang, J.; Zhu, L.; Shen, L. Comparison of Reference Gene Expression Stability in Mouse Skeletal Muscle via Five Algorithms. PeerJ 2022, 10, e14221. 10.7717/peerj.14221.

(66) Azhar, M.; Wardhani, B. W. K.; Renesteen, E. The Regenerative Potential of Pax3/Pax7 on Skeletal Muscle Injury. Journal of Genetic Engineering and Biotechnology 2022, 20 (1), 143. 10.1186/s43141-022-00429-x.

(67) Zammit, P. S.; Relaix, F.; Nagata, Y.; Ruiz, A. P.; Collins, C. A.; Partridge, T. A.; Beauchamp, J. R. Pax7 and Myogenic Progression in Skeletal Muscle Satellite Cells. Journal of Cell Science 2006, 119 (9), 1824–1832. 10.1242/jcs.02908.

(68) Yamamoto, M.; Legendre, N. P.; Biswas, A. A.; Lawton, A.; Yamamoto, S.; Tajbakhsh, S.; Kardon, G.; Goldhamer, D. J. Loss of MyoD and Myf5 in Skeletal Muscle Stem Cells Results in Altered Myogenic Programming and Failed Regeneration. Stem Cell Reports 2018, 10 (3), 956–969. 10.1016/j.stemcr.2018.01.027.

(69) Ganassi, M.; Badodi, S.; Wanders, K.; Zammit, P. S.; Hughes, S. M. Myogenin Is an Essential Regulator of Adult Myofibre Growth and Muscle Stem Cell Homeostasis. eLife 2020, 9, e60445. 10.7554/eLife.60445.

(70) C2C12 cells: biophysical, biochemical, and immunocytochemical properties - PubMed. https://pubmed.ncbi.nlm.nih.gov/8023908/ (accessed 2025-07-14).

(71) Nolan, A.; Heaton, R. A.; Adamova, P.; Cole, P.; Turton, N.; Gillham, S. H.; Owens, D. J.; Sexton, D. W. Fluorescent Characterization of Differentiated Myotubes Using Flow Cytometry. Cytometry A 2024, 105 (5), 332–344. 10.1002/cyto.a.24822.

(72) Shu, X. Z.; Ghosh, K.; Liu, Y.; Palumbo, F. S.; Luo, Y.; Clark, R. A.; Prestwich, G. D. Attachment and Spreading of Fibroblasts on an RGD Peptide–Modified Injectable Hyaluronan Hydrogel. Journal of Biomedical Materials Research Part A 2004, 68A (2), 365–375. 10.1002/jbm.a.20002.

(73) Alcazar, C. A.; Hu, C.; Rando, T. A.; Huang, N. F.; Nakayama, K. H. Transplantation of Insulin-like Growth Factor-1 Laden Scaffolds Combined with Exercise Promotes Neuroregeneration and Angiogenesis in a Preclinical Muscle Injury Model. Biomater. Sci. 2020, 8 (19), 5376–5389. 10.1039/D0BM00990C.

(74) Dreher, S. I.; Grubba, P.; Von Toerne, C.; Moruzzi, A.; Maurer, J.; Goj, T.; Birkenfeld, A. L.; Peter, A.; Loskill, P.; Hauck, S. M.; Weigert, C. IGF1 Promotes Human Myotube Differentiation toward a Mature Metabolic and Contractile Phenotype. American Journal of Physiology-Cell Physiology 2024, 326 (5), C1462–C1481. 10.1152/ajpcell.00654.2023.

(75) Pawlikowski, B.; Vogler, T. O.; Gadek, K.; Olwin, B. B. Regulation of Skeletal Muscle Stem Cells by Fibroblast Growth Factors. Developmental Dynamics 2017, 246 (5), 359–367. 10.1002/dvdy.24495.

(76) Wroblewski, O. M.; Vega-Soto, E. E.; Nguyen, M. H.; Cederna, P. S.; Larkin, L. M. Impact of Human Epidermal Growth Factor on Tissue-Engineered Skeletal Muscle Structure and Function. Tissue Engineering Part A 2021, 27 (17–18), 1151–1159. 10.1089/ten.tea.2020.0255.

(77) Wu, D.; Eugenis, I.; Hu, C.; Kim, S.; Kanugovi, A.; Yue, S.; Wheeler, J. R.; Fathali, I.; Feeley, S.; Shrager, J. B.; Huang, N. F.; Rando, T. A. Bioinstructive Scaffolds Enhance Stem Cell Engraftment for Functional Tissue Regeneration. Nat. Mater. 2025, 1–11. 10.1038/s41563-025-02212-y.

(78) Hao, L.; Yu, D.; Hou, X.; Zhao, Y. Advances in Polypyrrole Nanofiber Composites: Design, Synthesis, and Performance in Tissue Engineering. Materials (Basel) 2025, 18 (13), 2965. 10.3390/ma18132965.

(79) Dong, R.; Ma, P. X.; Guo, B. Conductive Biomaterials for Muscle Tissue Engineering. Biomaterials 2020, 229, 119584. 10.1016/j.biomaterials.2019.119584.

(80) Browe, D.; Freeman, J. Optimizing C2C12 Myoblast Differentiation Using Polycaprolactone-Polypyrrole Copolymer Scaffolds. J Biomed Mater Res A 2019, 107 (1), 220–231. 10.1002/jbm.a.36556.

(81) Taye, N.; Stanley, S.; Hubmacher, D. Stable Knockdown of Genes Encoding Extracellular Matrix Proteins in the C2C12 Myoblast Cell Line Using Small-Hairpin (Sh)RNA. J Vis Exp 2020, No. 156, 10.3791/60824.

(82) Willett, N. J.; Krishnan, L.; Li, M. T. A.; Guldberg, R. E.; Warren, G. L. Guidelines for Models of Skeletal Muscle Injury and Therapeutic Assessment. Cells Tissues Organs 2016, 202 (3–4), 214–226. 10.1159/000445345.

(83) Anderson, S. E.; Han, W. M.; Srinivasa, V.; Mohiuddin, M.; Ruehle, M. A.; Moon, J. Y.; Shin, E.; San Emeterio, C. L.; Ogle, M. E.; Botchwey, E. A.; Willett, N. J.; Jang, Y. C. Determination of a Critical Size Threshold for Volumetric Muscle Loss in the Mouse Quadriceps. Tissue Engineering Part C: Methods 2019, 25 (2), 59–70. 10.1089/ten.tec.2018.0324.

