## Supplemental Information for "Hyaluronic Acid-Coated Melt Electrowritten Scaffolds Promote Myoblast Attachment, Alignment, and Differentiation"

**Title:**

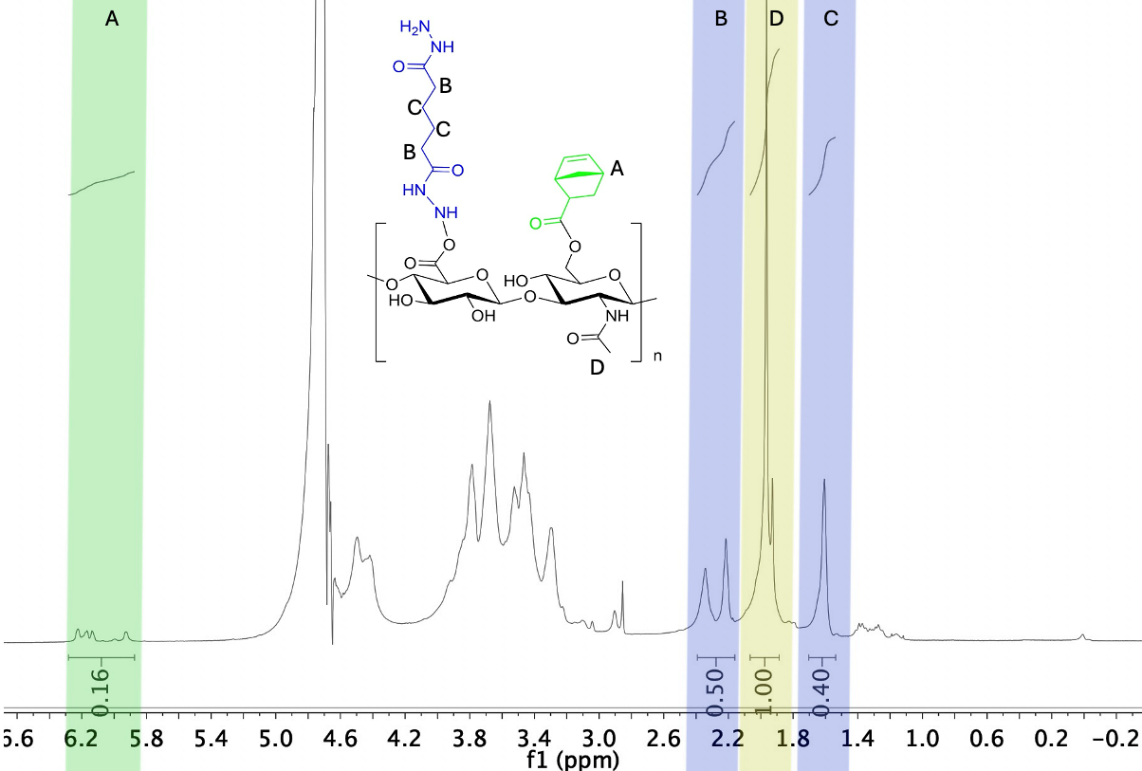

**Fig. S1** Representative ^1^H-NMR spectrum confirming modification of HA with norbornene and adipic acid dihydrazide functional groups.

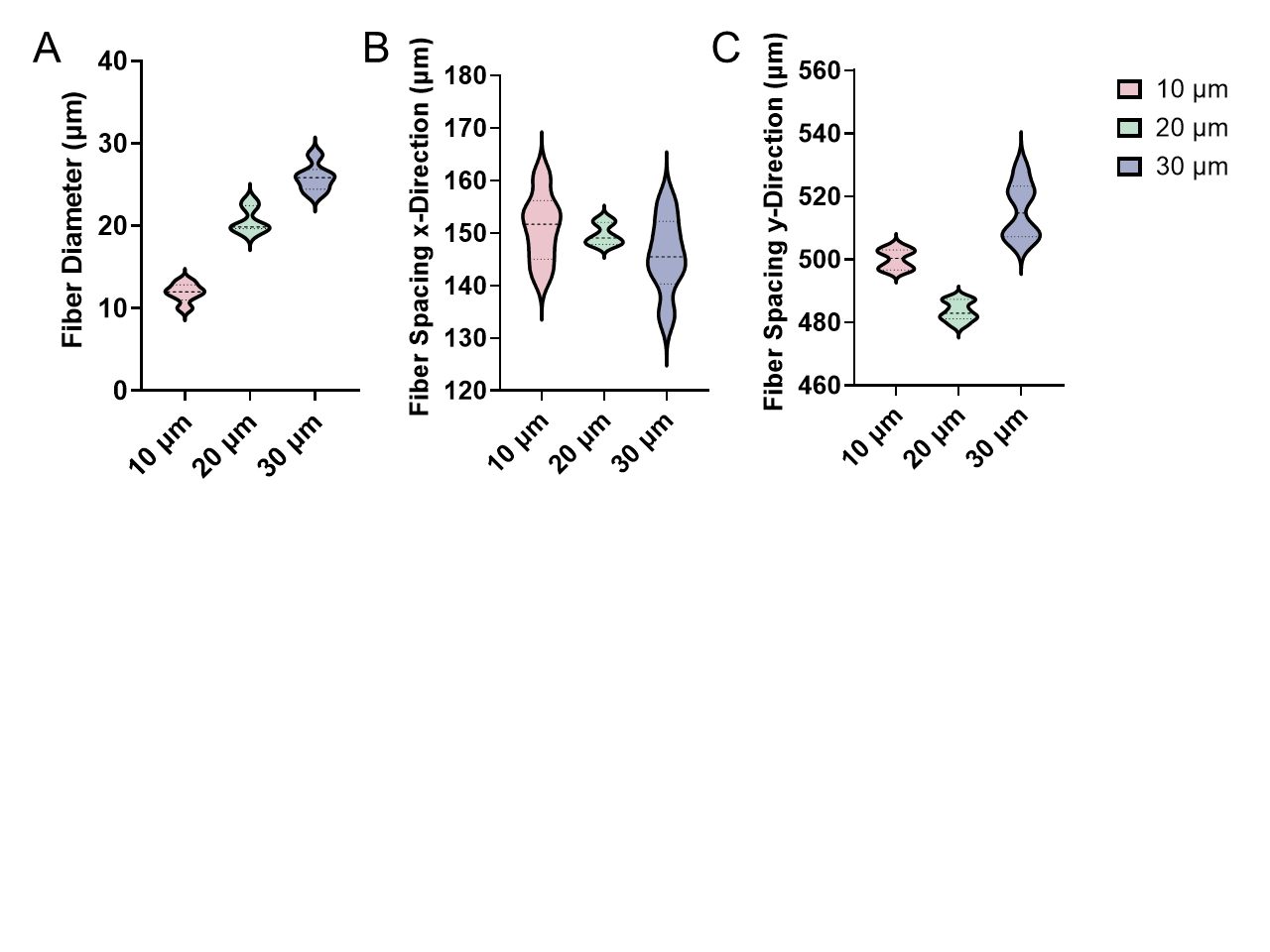

**Fig. S2** Measurements of MEW scaffold fiber diameters and fiber spacing using SEM images.

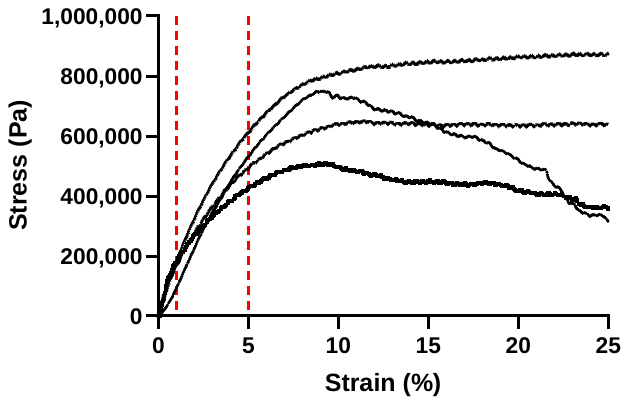

**Fig. S3** Stress-strain curves for determining the stiffness of 20 µm non-coated scaffolds. The slope of the linear region between 1-5% strain, indicated by dashed red lines, was used to determine the stiffness of the scaffolds.

| **Table S1. Summary of Primer Sequences** | | |
| --- | --- | --- |
|  | **Forward Sequence (3'-5')** | **Reverse Sequence (3'-5')** |
| **Pax3** | GCG TCT CTA AGA TCC TGT GCA G | GAT TTC CCA GCT AAA CAT GCC CG |
| **Pax7** | GAG TTC GAT TAG CCG AGT GC | CGG GTT CTG ATT CCA CAT CT |
| **Myf5** | AAC CAG AGA CTC AAG GT | GCT GGA CAA GCA ATC CAA GC |
| **MyoD** | AGT GAA TGA GGC CTT CGA GA | GCA TCT GAG TCG CCA CTG TA |
| **MyoG** | GAA GAA AAG GGA CTG GGG AC | GCG CAG GAT CTC CAC TTT AG |
| **GAPDH** | TCA ACG ACC CCT TCA TTG AC | ATG CAG GGA TGA TGT TCT GG |

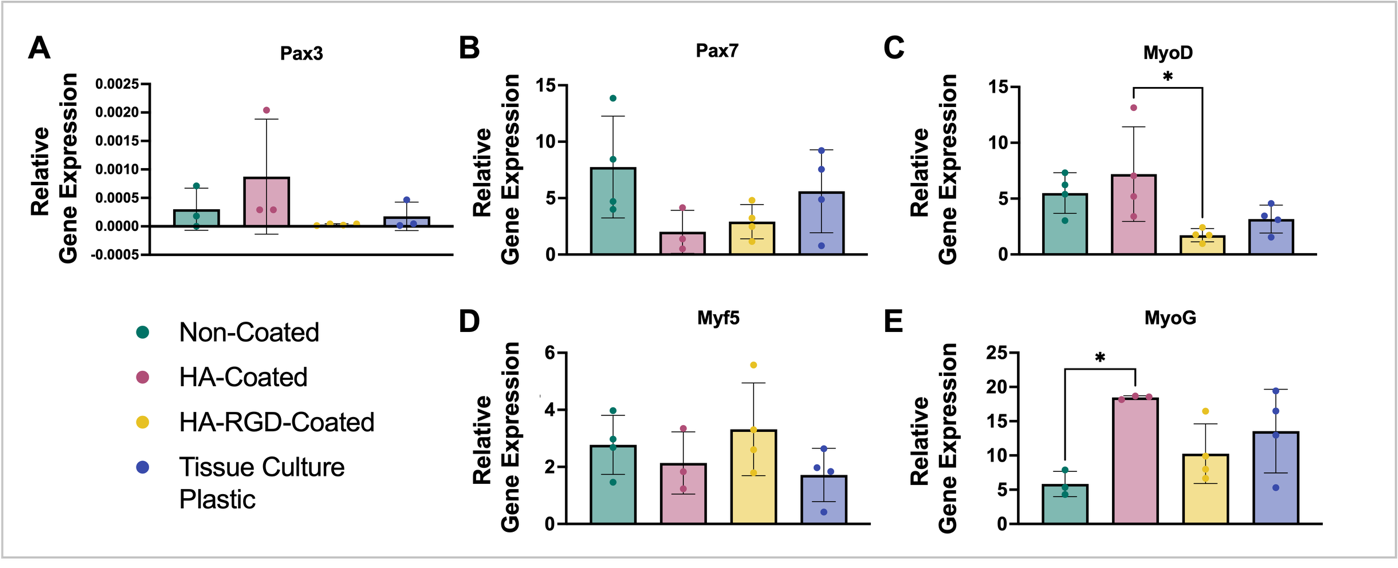

**Fig. S4** Myogenic gene expression of C2C12 myoblasts differentiated on non-coated, HA-coated, and HA-RGD-coated 20 µm fiber scaffolds or tissue culture plastic for 5 days. RT-qPCR was performed on differentiated cells and compared to undifferentiated cells and a GAPDH housekeeping gene to assess Pax3, Pax7, MyoD, Myf5, and MyoG gene expression, n=3-4. (One-way ANOVA with post-hoc Tukey's multiple comparisons test, * p < 0.05 as indicated)
